# Orthogonal synthetases for polyketide precursors

**DOI:** 10.1101/2022.02.28.482149

**Authors:** Riley Fricke, Cameron V. Swenson, Leah Tang Roe, Noah Hamlish, Omer Ad, Sarah Smaga, Christine L. Gee, Alanna Schepartz

**Author notes:** These authors contributed equally (Riley Fricke, Cameron V. Swenson).

## Abstract

The absence of orthogonal aminoacyl-tRNA synthetases that accept non-L-α-amino acids is the primary bottleneck hindering the *in vivo* translation of sequence-defined hetero-oligomers. Here we report PylRS enzymes that accept α-hydroxy acids, α-thio acids, *N*-formyl-L-α-amino acids, and α-carboxyl acid monomers (malonic acids) that are formally precursors to polyketide natural products. These monomers are all accommodated and accepted by the translation apparatus *in vitro*. High-resolution structural analysis of the complex between one such PylRS enzyme and a *meta*-substituted 2-benzylmalonate derivative reveals an active site that discriminates *pro*-chiral carboxylates and accommodates the large size and distinct electrostatics of an α-carboxyl acid substituent. This work emphasizes the potential of PylRS-derived enzymes for acylating tRNA with monomers whose α-substituent diverges significantly from the α-amine embodied in proteinogenic amino acids. These enzymes could act in synergy with natural or evolved ribosomes to generate diverse sequence-defined non-protein hetero-oligomers.

## Main

All extant organisms biosynthesize polypeptides in an mRNA template-dependent manner using a translation apparatus composed of ribosomes, aminoacyl-tRNA synthetases, tRNAs, and a host of ancillary factors. Repurposing this translation apparatus for the templated synthesis of mixed-sequence hetero-oligomers–especially those containing non-L-α-amino acids–would provide a biological route to sequence-defined non-peptide polymers with novel, tunable, evolvable properties and protein therapeutics with improved stability and expanded recognition potential. Chemical methods support the synthesis of certain sequence-defined (as opposed to sequence-controlled) non-peptide polymers^1,2^ but are limited to small scale and generate considerable waste. Cellular methods based on the translation apparatus generate less waste, especially on a large scale, achieve greater chain lengths, and are scalable for industrial production. Over the past decade, it has become clear that many non-L-α-amino acids, including α-hydroxy acids^3,4^, D-α-^5^, linear^6–8^ and cyclic^9^ β-, γ-^10,11^, and long chain amino acids^12^, α-aminoxy and α-hydrazino acids^13^, α-thio acids^14^, aramids^15,16^–even 1,3-dicarbonyl monomers that resemble polyketide precursors^16^–are accepted by *E. coli* ribosomes in small-scale *in vitro* reactions. The structural and electronic diversity of these monomers reiterates the important role of proximity in promoting bond-forming reactions within the *E. coli* peptidyl transferase center (PTC)^17^. However, there are scant examples in which non-L-α-amino acids have been incorporated into proteins *in vivo*^18–23^. The absence of orthogonal aminoacyl-tRNA synthetase (aaRS) enzymes that accept non-L-α-amino acid substrates is the primary bottleneck limiting the production of sequence-defined non-peptide polymers *in vivo* using wild-type or engineered ribosomes.

In *E. coli*, two families of orthogonal aaRS enzymes have been employed widely for the incorporation of hundreds of different non-canonical α-amino acid monomers^24–28^. The first includes pyrrolysyl-tRNA synthetase (PylRS) enzymes from methanogenic archaea and bacteria whose natural substrate is pyrrolysine^29^, an L-α-amino acid found in the active sites of certain enzymes involved in methane metabolism^30^. The second includes a large family of enzymes derived from *Methanocaldococcus jannaschii* tyrosyl-tRNA synthetase (*Mj*TyrRS)^24,31^. These two enzyme classes differ in how they recognize the α-amine of a bound substrate. While *Mj*TyrRS recognizes the substrate α-amine *via* multiple, direct hydrogen bonds to the side chains of Q173, Q176, and Y151 (PDB: 1J1U)^32^, *Methanosarcina mazei* PylRS (*Mm*PylRS) instead uses water-mediated hydrogen bonds to the N346 side chain and L301 and A302 backbone amides (PDB: 2ZCE; Fig. 1a)^33^. These differences were exploited by Kobayashi *et al*. to acylate the cognate tRNA *Mm*-tRNA^Pyl^ with a series of conservative *N*^ε^-Boc-L-lysine (L-BocK) analogs containing -OH, -H and -NHCH_3_ in place of the α-amine (Fig. 1b)^20^.

**Figure 1.**
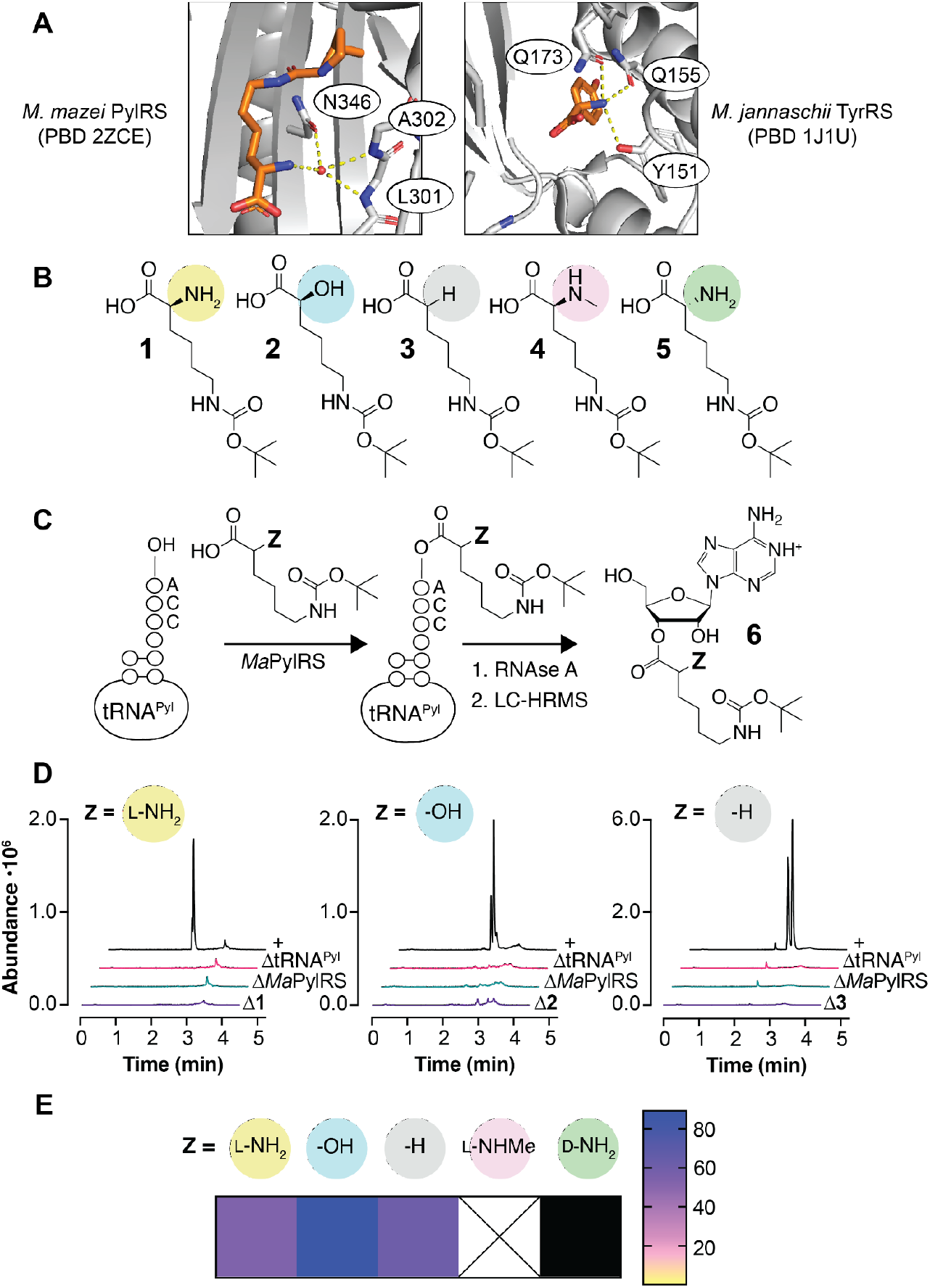
Promiscuous activity of wild-type *Ma*PylRS. **a**, The α-amines of L-α-amino acids are recognized differently by *M. mazei* PylRS (*Mm*PylRS, left)^34^ and *M. jannaschii* TyrRS (*Mj*TyrRS, right)^33^. **b**, *N*^ε^-Boc-L-lysine (L-BocK, **1**) analogs evaluated as substrates for *Ma*PylRS. **c**, Ribonuclease A (RNAse A) assay used to detect acylation of *Ma*-tRNA^Pyl^ with BocK analogs shown in panel B. **d**, LC-HRMS analysis of *Ma*-tRNA^Pyl^ acylation reactions after RNAse A digestion. Peak masses correspond to adenosine nucleoside **6** acylated on the 2’ or 3’ hydroxyl with the indicated monomer. **e**, Heat map illustrating the relative activities of substrates **1-5** for *Ma*PylRS as determined by intact tRNA analysis (Supplementary Fig. 3) as described in **Methods**. Reported yields are based on intact tRNA analysis. Black indicates no reaction product detected; X indicates the substrate was not investigated.

We hypothesized that the water-mediated α-amine recognition employed by *Mm*PylRS could provide sufficient space and flexibility for substrates with less conservative substituents in place of the α-amine. Here we report that the tolerance of *Mm*PylRS for substrates with multiple alternative substituents in place of the α-amine extends to *Methanomethylophilus alvus* PylRS (*Ma*PylRS)^34,35^ as well as *Ma*PylRS variants (*Ma*FRS1 and *Ma*FRS2)^36^ that recognize diverse L-phenylalanine derivatives. More importantly, *Ma*FRS1 and *Ma*FRS2 also accept phenylalanine derivatives with α-thio, *N*-formyl-L-α-amino, as well as an α-carboxyl substituent: 2-benzylmalonic acid. A final variant, *Ma*FRSA^37^, is selective for ring-substituted 2-benzylmalonate derivatives over L-Phe. Malonates contain a 1,3-dicarbonyl unit that represents the defining backbone element of polyketide natural products, and after decarboxylation have the potential to support Claisen-type condensation within the PTC to form a carbon-carbon bond. Structural analysis of *Ma*FRSA complexed with a *meta*-substituted 2-benzylmalonate derivative and a non-hydrolyzable ATP analogue reveals how the enzyme uses a novel pattern of hydrogen bonds to differentiate the two pro-chiral carboxylates in the substrate and accommodate the larger size and distinct electrostatics of an α-carboxyl substituent. *In vitro* translation studies confirm that tRNAs carrying α-thio, α-carboxyl, and *N*-formyl-L-α-amino acid monomers are effectively delivered to and accommodated by *E. coli* ribosomes. This work describes the first orthogonal aminoacyl-tRNA synthetase that accepts α-thio acids and the α-carboxyl acid precursors capable of effecting carbon-carbon bond formation within the ribosome. These novel activities emphasize the potential of PylRS as a scaffold for evolving new enzymes that act in synergy with natural or evolved ribosomes to generate diverse sequence-defined non-protein heterooligomers.

## Results

### *Ma*PylRS retains much of the promiscuity of *Mm*PylRS

First we set out to establish whether the novel substrate scope of *Mm*PylRS reported for L-BocK analogs (Fig. 1b)^20^ was retained by *Ma*PylRS, which offers advantages over *Mm*PylRS because it lacks the poorly soluble N-terminal tRNA-binding domain^38^ and is easier to express and evaluate *in vitro*^35^. The C-terminal catalytic domain of *Mm*PvlRS^33^ is 36% identical to *Ma*PylRS and the structures are largely superimposable (Supplementary Fig. 1a,b). To evaluate whether D-BocK as well as L-BocK analogs with -OH or -H in place of the α-amino group were substrates for *Ma*PylRS, we made use of a validated RNAse A/LC-HRMS assay^16,39^. This assay exploits RNAse A to cleave the phosphodiester bond of unpaired C and U residues to generate 2’, 3’-cyclic phosphate products^40^. As a result, the residue at the tRNA 3’ terminus is the only mononucleoside product lacking a phosphate (Fig. 1c). Incubation of L-BocK **1** (2 mM) with purified *Ma*PylRS (2.5 μM) and *Ma*-tRNA^Pyl^ (25 μM) (Supplementary Fig. 2) at 37 °C for 2 hours led to a pair of RNAse A digestion products whose expected mass (496.25142 Da) corresponds to the adenosine nucleoside **6** as a mixture of 2’- and 3’-acylated species (Fig. 1d)^41^. No products with this mass were observed when the reaction mixture lacked *Ma*-tRNA^Pyl^, L-BocK, or *Ma*PylRS, and mass analysis of the intact tRNA product confirmed a 53% yield of acylated tRNA (**1-tRNA^Pyl^**, Supplementary Fig. 3a). Under these conditions, L-BocK analogs with either -OH (**2**) or -H (**3**) in place of the α-amino group were also substrates for *Ma*PylRS as judged by RNAse A (Fig. 1d) and intact tRNA mass spectrometry assays with acylated tRNA yields of 89% (**2-tRNA^Pyl^**) and 61% (**3-tRNA^Pyl^**) (Supplementary Fig. 3b,c, respectively). No reactivity was detected with D-BocK (Supplementary Figs. 3d and 7). We conclude that *Ma*PylRS retains much (although not all) of the previously reported^20^ promiscuity of *Mm*PylRS. We note that certain non-natural monomers with relatively high activity, including α-hydroxy **2** and des-amino **3**, led to measurable levels (2-13%) of diacylated tRNA (Supplementary Table 3). Diacylated tRNAs have been observed as products in cognate reactions of *T. thermophilus* PheRS^42^ and are active in prokaryotic translation^43^.

### *Ma*PylRS variants retain activity for phenylalanine derivatives with diverse α-amine substitutions

PylRS is a subclass IIc aaRS that evolved from PheRS^44^*. Mm*PylRS variants with substitutions at two positions that alter the architecture of the side chain-binding pocket (N346, C348) (Supplementary Fig. 1a) accept L-phenylalanine and its derivatives in place of pyrrolysine^36,44,25^. We integrated the mutations in two variants that accept unsubstituted L-Phe (*Mm*FRS1 and *Mm*FRS2)^36^ into the *Ma*PylRS sequence to generate *Ma*FRS1 (N166A, V168L) and *MaFRS2* (N166A, V168K) (Supplementary Fig. 2). We then used RNAse A and intact tRNA mass spectrometry assays to determine if *Ma*FRS1 or *Ma*FRS2 retained activity for L-phenylalanine **7** and analogs in which the L-α-amino group was substituted by -OH **8**, -H **9**, -NHCH_3_ **10,** or D-NH_2_ **11** (Fig. 2a).

**Figure 2.**
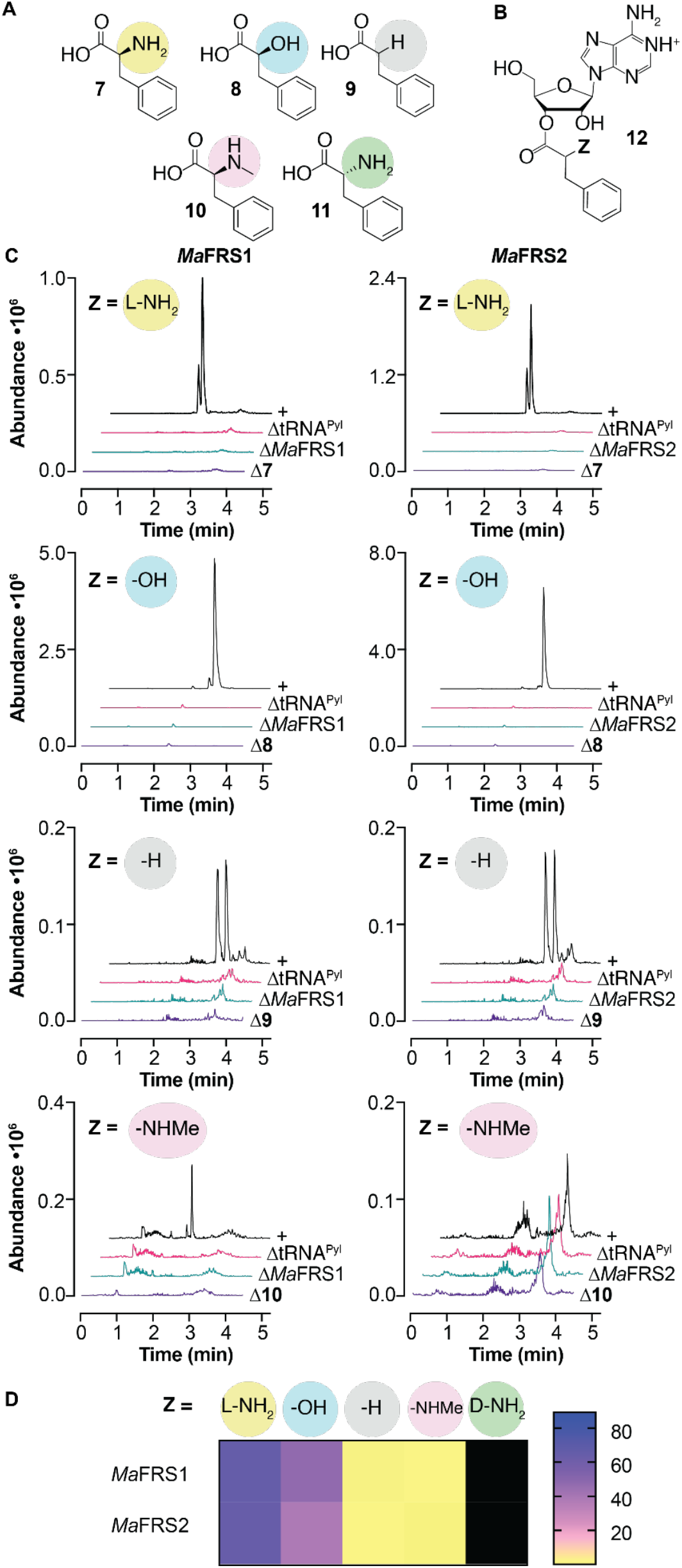
*Ma*FRS1 and *Ma*FRS2 process phenylalanine analogs with substitutions at the α-amine. **a**, Phenylalanine analogs evaluated as substrates for *Ma*FRS1 and *Ma*FRS2. **b**, Adenosine nucleoside formed during RNAse A digestion of acyl-tRNA. **c**, LC-HRMS analysis of *Ma*-tRNA^Pyl^ acylation reactions after RNAse A digestion. Adenine nucleoside **12** acylated on the 2’- or 3’-hydroxyl of the 3’ terminal ribose of *Ma*-tRNA^Pyl^ could be detected in *Ma*FRS1 and *Ma*FRS2 reactions with L-Phe **7** and substrate **8**; substrates **9** and **10** (Z = -H, -NHCH_3_) showed more modest reactivity. **d**, Heat map illustrating the relative yields with substrates **7**-**11** for *Ma*FRS1 and *Ma*FRS2 as determined by intact tRNA analysis (Supplementary Figs. 4 and 5) as described in **Methods**. Black indicates no reaction product detected.

Incubation of L-Phe **7** (10 mM) with purified *Ma*FRS1 or *Ma*FRS2 (2.5 μM) and *Ma*-tRNA^Pyl^ (25 μM) at 37 °C for 2 hours led in both cases to a pair of products whose expected mass (415.17244 Da) corresponds to adenosine nucleoside **12** (Z = L-NH_2_, Fig. 2b) as the expected mixture of 2’- and 3’-acylated products (Fig. 2c). No product with this mass was observed when the reaction mixture lacked *Ma*-tRNA^Pyl^, L-Phe, or *Ma*FRS1 or *Ma*FRS2, and mass analysis of the intact tRNA product confirmed a 66% (*Ma*FRS1) and 65% (*Ma*FRS2) yield of acylated tRNA (**7-tRNA^Pyl^**) (Supplementary Figs. 4a and 5a, respectively). L-Phe analogs **8-10** were all substrates for both *Ma*FRS1 and *Ma*FRS2 as judged by both RNAse A (Fig. 2c) and intact tRNA analysis (Supplementary Figs. 4a-d and 5a-d), with reactivities in the order L-α-amino **7** > α-hydroxy **8** >> des-amino **9** ~ *N*-Me-L-α-amino **10** based on intact tRNA yields (Fig. 2d). Mono- and di-acylated tRNA products were observed for substrates **7** and **8** (Supplementary Table 3).

Interestingly, despite the fact that the des-amino-BocK analog **3** was a strong substrate for the wild-type *Ma*PylRS, the des-amino-Phe analog **9** had only modest activity with *Ma*FRS1 and *Ma*FRS2, as observed in the RNAse assay (Fig. 2c and Supplementary Figs. 4c and 5c). Again, no reactivity was detected with D-Phe (Supplementary Figs. 4e, 5e, and 7).

### *Ma*FRS1 & *MaFRS2* process substrates with novel α-substituents

We then began to explore phenylalanine analogs with larger and electrostatically distinct functional groups at the α-carbon: α-thio acids, *N*-formyl-L-α-amino acids, and α-carboxyl acids (malonates) (Fig. 3a). α-thio acids are substrates for extant *E. coli* ribosomes in analytical-scale *in vitro* translation reactions with yields as high as 87% of the corresponding α-amino acids^14^, and thioesters can have a half-life in *E. coli* of more than 36 hours^45^. Peptides and proteins containing thioesters could also act as substrates for PKS modules^46^ to generate unique keto-peptide natural products. Formylation of methionyl-tRNA is important for initiation complex formation, and formylation could enhance initiation with non-methionyl α-amino acids *in vivo*^47,48^. Moreover, *E. coli* ribosomes incorporate monomers containing a 1,3-dicarbonyl moiety at the peptide N-terminus to produce keto-peptide hybrids^16^. To our knowledge, there are currently no aaRS enzymes, orthogonal or otherwise, that accept α-thio, *N*-formyl-L-α-amino, or α-carboxyl acid substrates to generate the acylated tRNAs required for *in vivo* translation (when extant ribosomes are compatible) or ribosome evolution (when extant ribosomes are incompatible).

**Figure 3.**
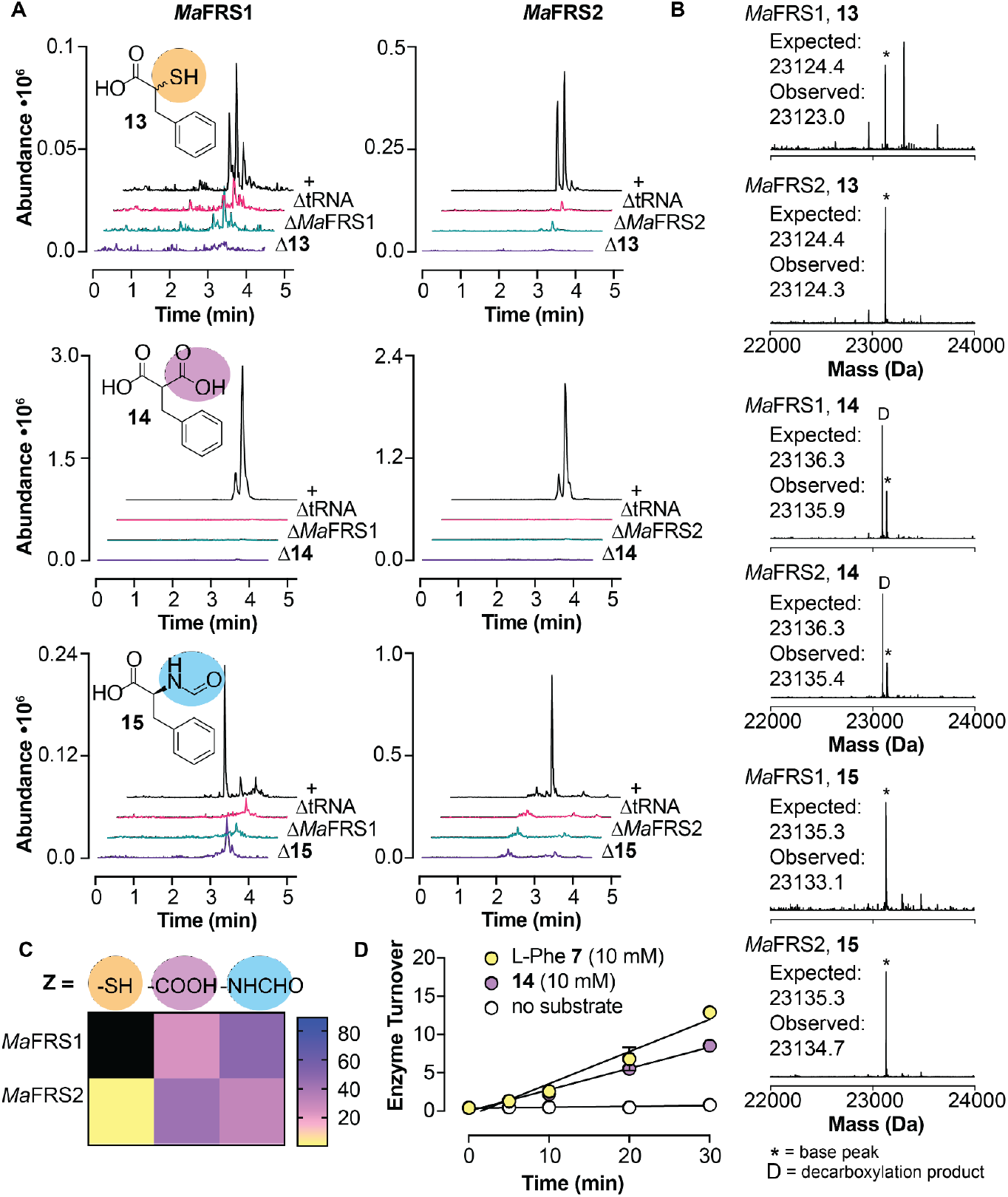
*Ma*FRS1 and *Ma*FRS2 process substrates bearing novel α-substituents. **a**, LC-HRMS analysis of *Ma*-tRNA^Pyl^ acylation reactions using *Ma*FRS1 or *Ma*FRS2 following RNAse A digestion. Adenosine nucleoside **12** acylated on the 2’- or 3’-hydroxyl of the 3’ terminal ribose of *Ma*-tRNA^Pyl^ could be detected in *Ma*FRS1 and *Ma*FRS2 reactions with α-thio acid **13**, α-carboxyl acid **14**, and *N*-formyl-L-Phe **15**. **b**, LC-MS analysis of intact tRNA products confirms that monomers **13**-**15** are substrates for *Ma*FRS1 and *Ma*FRS2. We note that intact tRNAs acylated with 2-benzylmalonate **14** showed evidence of decarboxylation (indicated by a D). No evidence for decarboxylation was observed when the same acyl-tRNAs were evaluated using the RNAse A assay (Supplementary Fig. 8), suggesting that decarboxylation occurs either during workup or during the LC-MS itself. **c**, Heat map illustrating the relative activities of substrates **13**-**15** with *Ma*FRS1 and *Ma*FRS2 as determined by intact tRNA analysis (Supplementary Figs. 4 and 5) as described in **Methods**. Black indicates no reaction product detected. Initially no acyl-tRNA was detected when *Ma*FRS1 was incubated with **13**, but when the enzyme concentration was increased 5-fold, acyl-tRNA was detected with mono- and diacyl yields of 0.7 and 9.7%, respectively. **d**, Turnover of *Ma*FRS1 over time with L-Phe **7** and 2-benzylmalonate **14** using the malachite green assay.

We found that L-Phe analogs in which the α-amine was substituted with α-thiol (**13)**, α-carboxyl (**14**), or *N*-formyl-L-α-amine (**15**) moieties were all substrates for *Ma*FRS1 and *Ma*FRS2 (Fig. 3a-c). In particular, α-carboxyl **14** and *N*-formyl-L-α-amine **15** were excellent substrates. Incubation of α-carboxyl acid **14** (10 mM) with *Ma*FRS1 or *Ma*FRS2 (2.5 μM) and *Ma*-tRNA^Pyl^ (25 μM) at 37 °C for 2 hours followed by digestion by RNAse A led to formation of a pair of products whose expected mass (444.15137 Da) corresponds to the adenosine nucleoside **12** (Z = -COOH). LC-MS analysis of intact tRNA products confirmed a 24% (*Ma*FRS1) and 43% (*Ma*FRS2) yield of acylated tRNA (**14-tRNA^Pyl^**) (Supplementary Figs. 4h and 5g, respectively). Because the α-carbon of substrate **14** is prochiral, mono-acylation of *Ma*-tRNA^Pyl^ can generate two diastereomeric product pairs–one pair in which the 3’-hydroxyl group is acylated by the *pro*-S or *pro*-R carboxylate and another in which the 2’-hydroxyl group is acylated by the *pro*-S or *pro*-R carboxylate. These diastereomeric products result from alternative substrate orientations within the enzyme active site (*vide infra*).

Incubation of *N*-formyl-L-α-amine (**15**) under identical conditions led to the expected adenosine nucleoside **12** (Z = -NHCHO) and LC-MS analysis confirmed a 49% (*Ma*FRS1) and 31% (*Ma*FRS2) yield of acylated tRNA (**15-tRNA^Pyl^**) (Supplementary Figs. 4i and 5h, respectively). The α-thio L-Phe analog **13** was also a substrate for *Ma*FRS1 and *Ma*FRS2, though higher concentrations of *Ma*FRS1 were necessary to observe the acyl-tRNA product (**13-tRNA^Pyl^**) using intact tRNA LC-MS (Supplementary Figs. 4f-g and 5f). These results illustrate that the active site pocket that engages the α-amine in PylRS can accommodate substituents with significant differences in mass (-NHCHO) and charge (-COO^-^). Despite these differences, 2-benzylmalonate **14** (2-BMA) is an excellent substrate for *Ma*FRS1 and *Ma*FRS2; kinetic analysis using the malachite green assay^49^ revealed a rate of adenylation by *Ma*FRS1 that was 66% of the rate observed for L-Phe (Fig. 3d). The tolerance for malonate substrates extends to WT *Ma*PylRS itself: the α-carboxylate analog of L-BocK **16** was also a measurable substrate for WT *Ma*PylRS (Supplementary Figs. 3e and 7).

### *Ma*FRSA processes *meta*-substituted 2-benzylmalonic acid substrates and is orthogonal to L-Phe

While *Ma*FRS1 and *Ma*FRS2 demonstrated the ability to process substrates with unusual α-substituents, they also process L-Phe with comparable efficiency (Fig. 2d), which would interfere with the selective charging of the non-L-α-amino acid. Variants of *Mm*PylRS that accept *para*-, *ortho*-, and *meta*-substituted L-Phe derivatives have been reported^37,50–53^. In particular, *Mm*PylRS containing two active site mutations (N346A and C348A; henceforth referred to as FRSA) shows high activity for L-Phe analogs with bulky alkyl substituents and low activity towards L-Phe^37^. We expressed and purified a variant of *Ma*PylRS containing these mutations (*Ma*FRSA: N166A, V168A) (Supplementary Fig. 2) and demonstrated that it shows high activity for derivatives of malonate **14** carrying *meta*-CH_3_ (**17**), *meta*-CF_3_ (**18**), and *meta*-Br (**19**) substituents and low activity for L-Phe using both RNAse A (Fig. 4a) and intact tRNA analysis (Supplementary Fig. 6a-d). Of the *meta*-substituted 2-benzylmalonates, *Ma*FRSA shows the highest activity for *meta*-CF_3_-2-benzylmalonate **18** (*meta*-CF_3_-2-BMA). Kinetic analysis^49^ revealed a rate of adenylation that was 36% of the rate observed for the L-α-amino acid counterpart *meta*-CF_3_-L-Phe (Fig. 4d). Although derivatives of malonate **14** carrying *meta*-CH_3_ (**17**), *meta*-CF_3_ (**18**), and *meta*-Br (**19**) substituents are also excellent substrates for *Ma*FRS1 (Supplementary Fig. 4j-l) and *Ma*FRS2 (Supplementary Fig. 5i-k), *Ma*FRSA has significantly lower activity for L-Phe, allowing for the selective acylation of tRNA with *meta*-substituted 2-benzylmalonates without interference from L-Phe (Fig. 4c). Indeed, when *Ma*FRSA is incubated with an equal concentration (10 mM) of L-Phe **7** and *meta*-CF_3_-2-BMA **18**, only the malonyl product (**18**-**tRNA^Pyl^**) is observed (Supplementary Fig. 6e).

**Figure 4.**
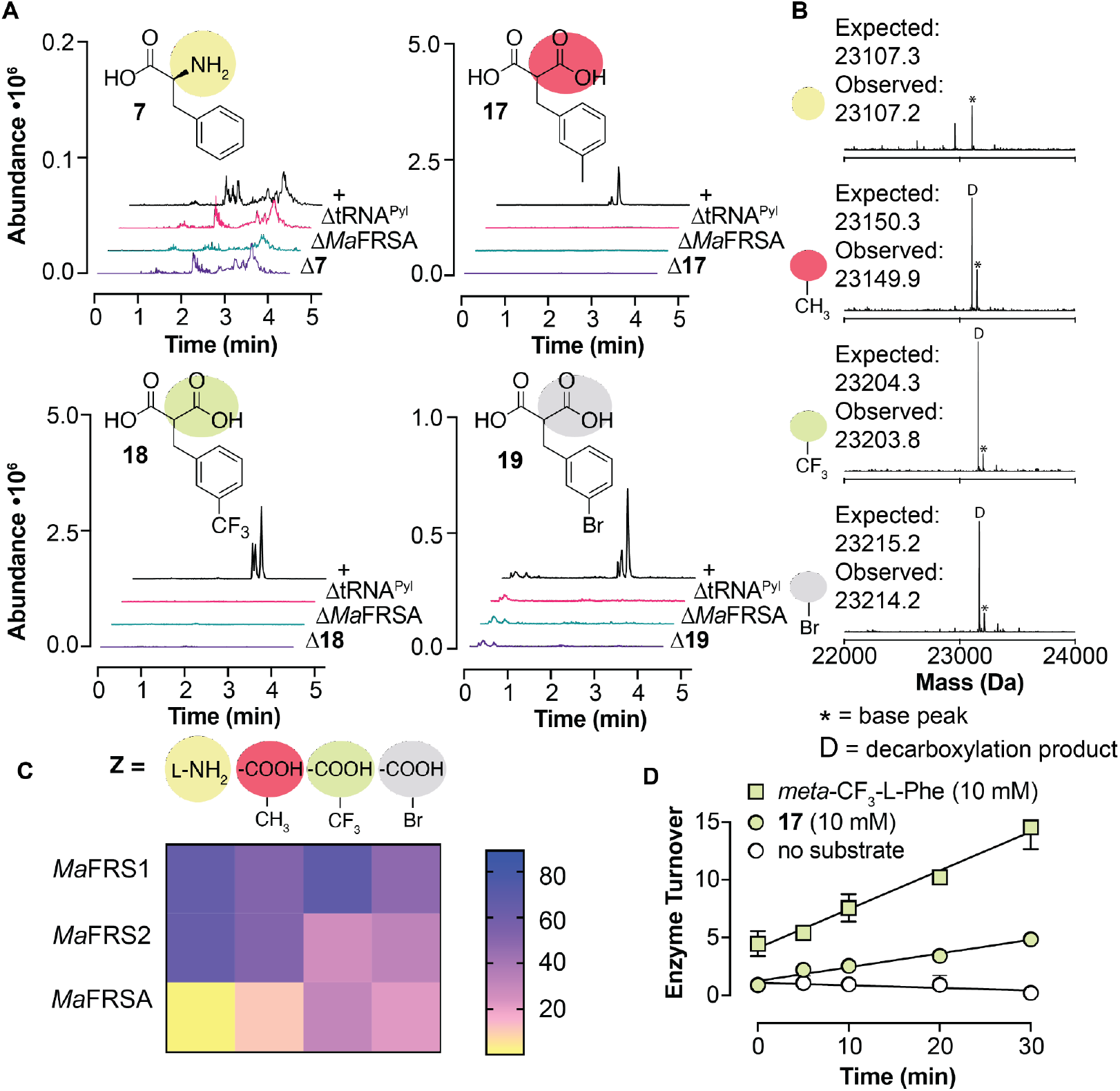
*Ma*FRSA selectively acylates *Ma*-tRNA^Pyl^ with *meta*-substituted 2-benzylmalonate derivatives. **a**, LC-HRMS analysis of *Ma*-tRNA^Pyl^ acylation products after digestion with RNAse A. **b**, LC-MS analysis of intact tRNA products confirms that *meta-substituted* 2-benzylmalonates **17**-**19** are substrates for *Ma*FRSA. We note that intact tRNAs acylated with *meta*-substituted 2-benzylmalonates **17**-**19** showed evidence of decarboxylation (indicated by a D). No evidence for decarboxylation was observed when the same acyl-tRNAs were evaluated using the RNAse A assay (Supplementary Fig. 8), suggesting that decarboxylation occurs either during workup or during the LC-MS itself. **c**, Heat map illustrating the relative activities of L-Phe **7** and substrates **17**-**19** with *Ma*FRS1, *Ma*FRS2, and *Ma*FRSA. Black indicates no reaction product detected. **d**, Turnover of *Ma*FRSA over time with *meta*-CF_3_-L-Phe and *meta*-CF_3_-2-BMA **18** using the malachite green assay.

### Structural analysis of *Ma*FRSA-*meta*-CF_3_-2-benzylmalonate complex reveals novel interactions

To better understand how 2-benzylmalonate substrates are accommodated by the *Ma*FRSA active site, we solved the crystal structure of *Ma*FRSA bound to both *meta*-CF_3_-2-BMA and the non-hydrolyzable ATP analog adenosine 5’-(β,γ-imido)triphosphate (AMP-PNP). The structure was refined at 1.8 Å with clear substrate density for *meta*-CF_3_-2-BMA visible at *1σ* in the *2Fo-Fc* map (Supplementary Fig. 9a-d). *Ma*FRSA crystallized with two protein chains in the asymmetric unit and an overall architecture resembling published PylRS structures (Fig. 5a)^33,54,55^. The two protein chains in the asymmetric unit are not identical and interact with different orientations of *meta*-CF_3_-2-BMA (Fig. 5b). One orientation of *meta*-CF_3_-2-BMA (chain A, light purple) mimics that of L-pyrrolysine (Pyl) bound to *Mm*PylRS ^33^ and would result in adenylation of the *pro*-R carboxylate (Fig. 5c); the other orientation (chain B, dark purple) would result in adenylation of the *pro*-S carboxylate (Fig. 5d).

**Figure 5.**
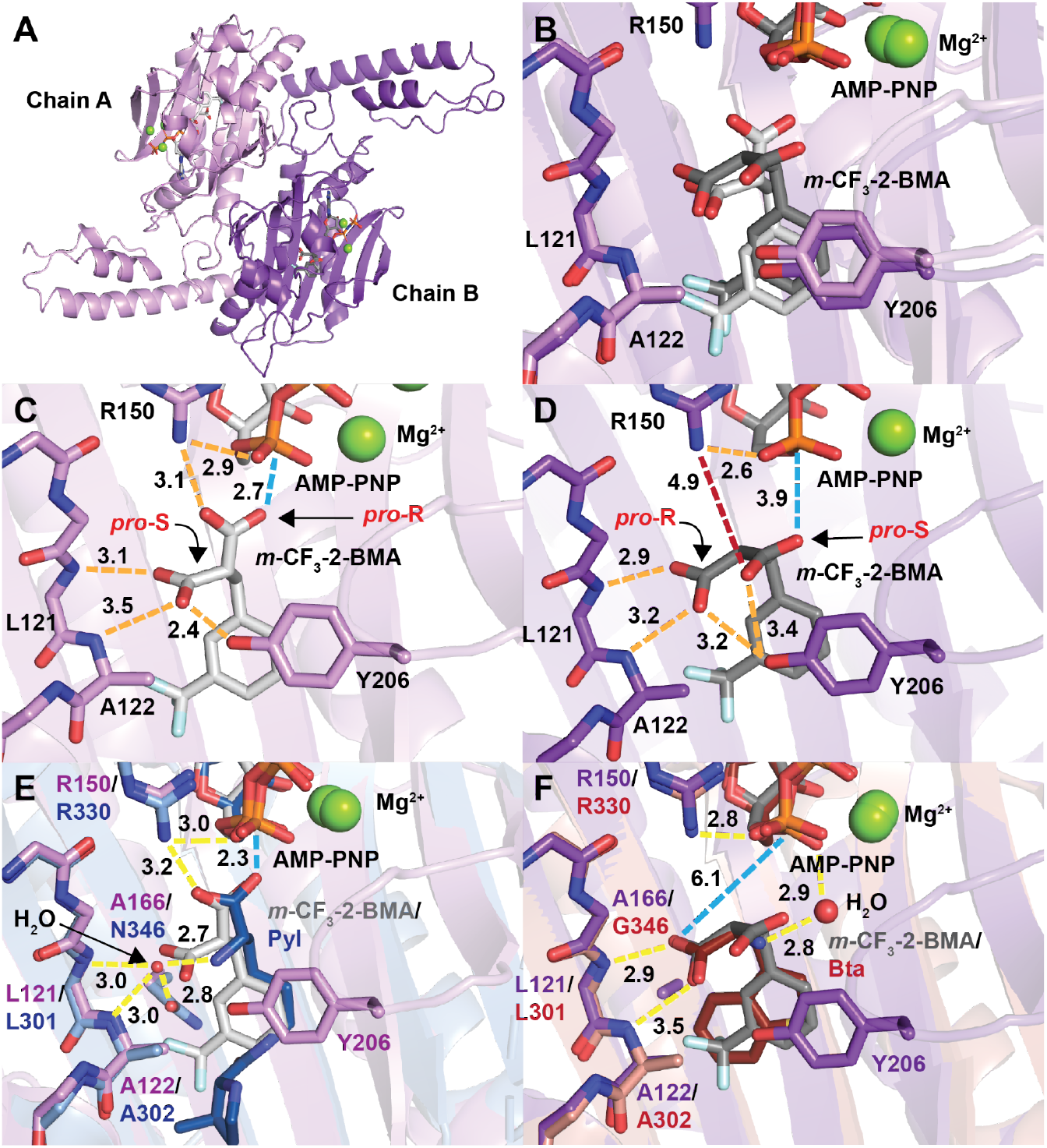
Structure of *Ma*FRSA bound to *meta*-CF_3_-2-BMA and AMP-PNP reveals basis for distinct reactivity at *pro*-R and *pro*-S substrate carboxylates. **a**, *Ma*FRSA dimer containing two nonidentical chains in the asymmetric unit. **b**, Alignment of the active sites of chains A (light purple) and B (dark purple) reveals *meta*-CF_3_-2-BMA (grays) bound in two alternate conformations. **c**, In chain A, *meta*-CF_3_-2-BMA is coordinated by an extensive hydrogen bond network (orange dashes) that positions the *pro*-R carboxylate oxygen for nucleophilic attack (blue dashes); interatomic distances are shown over dashed lines in Å. **d**, In chain B, *meta-CF_3_-2-BMA* is coordinated by similar hydrogen bonds, but in this case the *pro*-S carboxylate is rotated away from AMP-PNP with a loss of the hydrogen bond to R150 (red dashes) and a longer distance between the *pro*-S carboxylate nucleophile and the α-phosphate of AMP-PNP. **e**, Alignment of active site A with WT *Mm*PylRS bound to Pyl and AMP-PNP (PDB: 2ZCE, blue)^34^ illustrates the difference between the water-mediated hydrogen bonds (yellow dashes) to the α-amine of Pyl in PylRS *versus* the direct carboxyl to backbone hydrogen bonding of *m*-CF_3_-2-BMA bound to *Ma*FRSA. **f**, Comparison of active site B with *Mm*BtaRS (N346G/C348Q) bound to Bta (PDB: 4ZIB, red)^57^ reveals similar binding modes with hydrogen bonds (yellow dashes) between the substrate carboxylate and amide backbone of the enzyme when N166/N346 (*Ma*/*Mm* numbering) is mutated.

Regardless of orientation, discrete networks of hydrogen bonds are used by *Ma*FRSA to discriminate between the *pro*-R and *pro*-S carboxylates of *meta*-CF_3_-2-BMA. In chain A, the *pro*-S carboxylate accepts a hydrogen bond from the backbone amides of L121 and A122 and the phenolic -OH of Y206 (Fig. 5c and Supplementary Fig. 9e). A hydrogen bond from the R150 guanidinium positions the *pro*-R carboxylate for nucleophilic attack with a distance of 2.7 Å between the carboxyl oxygen and the α-phosphorous of AMP-PNP. In chain B, the orientation of *meta*-CF_3_-2-BMA is flipped relative to the conformation bound to chain A (Fig. 5d and Supplementary Fig. 9f). The *pro*-R carboxylate accepts a hydrogen bond from the backbone amides of L121 and A122 and the phenolic -OH of Y206 as seen for the *pro*-S carboxylate in chain A. However, in chain B the *pro*-S carboxylate is rotated away from AMP-PNP and towards Y206 resulting in loss of the hydrogen bond to R150 and a longer distance of 3.9 Å between the carboxyl oxygen and the α-phosphorous of AMP-PNP. RNAse A analysis of *Ma*-tRNA^Pyl^ acylation by *meta*-CF_3_-2-BMA shows more than two peaks of identical mass (Fig. 4a) that likely correspond to the two diastereomeric pairs formed from attack of the 2’- or 3’ - tRNA hydroxyl group on the activated *pro*-R or *pro*-S carboxylate. More than two peaks with identical mass are also observed as RNAse A digestion products in *Ma*-tRNA^Pyl^ acylation reactions of other *meta*-substituted 2-benzylmalonates (Fig. 4a). While the *meta*-CF_3_-2-BMA conformation in chain A appears more favorably positioned for catalysis, suggesting that the *pro*-R carboxylate is acylated preferentially, the appearance of more than two peaks in the RNAse A assay suggests that both conformations are catalytically competent.

As anticipated, the non-reactive, *pro*-S carboxylate of *meta*-CF_3_-2-BMA is recognized by *Ma*FRSA chain A using interactions that are distinct from those used by *Mm*PylRS to recognize the Pyl α-amine. In the *Mm*PylRS:Pyl:AMP-PNP complex (PDB: 2ZCE)^33^, the Pyl α-amine is recognized by water-mediated hydrogen bonds to the backbone amides of L301 and A302 and the side chain carbonyl of N346, rather than by direct hydrogen bonds to the backbone amides of L121 and A122 as seen for recognition of the non-reacting carboxylate of *meta*-CF_3_-2-BMA by both chains of *Ma*FRSA (Fig. 5e). The interactions used to recognize the non-reacting carboxylate of *meta*-CF_3_-2-BMA are, however, similar to those used to recognize the single carboxylate of diverse α-amino acids by other PylRS variants with mutations at N166/N346 (*Ma*/*Mm* numbering). For example, the structures of *Mm*PylRS variants with mutations at N346, such as *Mm*IFRS and *Mm*BtaRS bound to 3-iodo-L-phenylalanine (3-I-F, PDB: 4TQD)^52^ and 3-benzothienyl-L-alanine (Bta, PDB: 4ZIB)^56^, respectively, show the substrate bound with the carboxylate directly hydrogen bonded to the L301 and A302 backbone amides, just as seen for *meta*-CF_3_-2-BMA bound to *Ma*FRSA (Fig. 5f and Supplementary Fig. 10a). In these cases, the bound water seen in the *Mm*PylRS:Pyl:AMP-PNP complex is either absent or displaced. Mutation of N166/N346 may destabilize the water-mediated hydrogen bonding between the substrate α-amine and backbone amides seen in wild-type PylRS and promote alternative direct hydrogen bonding of a substrate carboxylate to backbone amides as seen in *Ma*FRSA, *Mm*IFRS, and *Mm*BtaRS.

The largest differences between *Ma*FRSA and other reported PylRS structures are localized to a 6-residue loop that straddles β-strands β7 and β8 and contains the active site residue Y206. In the the *Mm*PylRS:Pyl:AMP-PNP structure (PDB: 2ZCE)^33^ and the wild-type *Ma*PylRS apo structure (PDB: 6JP2)^54^, the β7-β8 loop is either unstructured or in an open conformation positioning Y206/Y384 away from the active site, respectively (Supplementary Fig. 10b). Among wild-type PylRS structures, the β7-β8 loop exists in the closed conformation only in the structure of *Mm*PylRS bound to the reaction product, Pyl-adenylate (PDB: 2Q7H)^55^. In this structure, Y384 accepts and donates a hydrogen bond to the Pyl-adenylate α-amine and pyrrole nitrogen, respectively, and forms a hydrophobic lid on the active site. In both chains of substrate-bound *Ma*FRSA, the non-reacting carboxylate of *meta*-CF_3_-2-BMA forms similar hydrogen bonds to Y206. These hydrogen bonds effectively close the β7-β8 loop to form a hydrophobic lid on the active site, which may contribute to the high acylation activity observed for *meta*-CF_3_-2-BMA with *Ma*FRSA. We note that the β7-β8 loop in chain A exhibits lower B-factors than in chain B indicating tighter binding and providing further evidence that chain A represents the more active binding mode of *meta*-CF_3_-2-BMA (Supplementary Fig. 10c,d). The two alternative poses of *meta*-CF_3_-2-BMA in chains A and B correspond to a ~120° degree rotation about the Cα-Cβ bond of the substrate that largely maintains interactions with the *meta*-CF_3_-2-BMA side chain but alters the placement of the reacting carboxylate. This observation emphasizes the dominant stabilizing role of the main chain H-bonds provided by the backbone amides of L121 and A122.

### *In vitro* translation initiation with novel monomers *via* codon skipping

We performed *in vitro* translation experiments to verify that the uniquely acylated derivatives of *Ma*-tRNA^Pyl^ produced using *Ma*PylRS variants are effectively shuttled to and accommodated by the *E. coli* ribosome. Whereas the *E. coli* initiator tRNA^fMet^ has been engineered into a substrate for *Mj*TvrRS variants to introduce non-canonical L-α-amino acids at the protein N-terminus^48^, *Ma*-tRNA^Pyl^ lacks the key sequence elements for recognition by *E. coli* initiation factors precluding its use for initiation *in vivo*^35,57^. It has been reported^58^ that in the absence of methionine, *in vitro* translation can begin at the second codon of the mRNA template, a phenomenon we refer to as “codon skipping”. We took advantage of codon skipping and a commercial *in vitro* translation (IVT) kit to evaluate whether *Ma*-tRNA^Pyl^ enzymatically charged with monomers **13**-**15** would support translation initiation.

To avoid competition with release factor 1 (RF1) at UAG codons, the anticodon of *Ma*-tRNA^Pyl^ was mutated to ACC (*Ma*-tRNA^Pyl^-ACC) to recode a glycine GGT codon at position 2 in the mRNA template. To maximize yields of acyl-tRNA, we increased the aaRS:tRNA ratio to 1:2 (monomers **7**, **14**, and **15**) or 1:1 (monomer **13**), extended the incubation time to 3 hours, and used the most active *Ma*FRSx variant for each monomer. These modifications resulted in acyl-tRNA yields of 79% (**7**, *Ma*FRS1), 13% (**13**, *Ma*FRS2), 85% (**14**, *Ma*FRS2), and 82% (**15**, *Ma*FRS1). The acylated *Ma*-tRNA^Pyl^-ACC was added with a DNA template encoding a short MGV-FLAG peptide (MGVDYKDDDDK) (Fig. 6a) to a commercial *in vitro* translation kit (PURExpress^®^ Δ (aa, tRNA), NEB). When methionine was excluded from the reaction mixture, translation initiated at the second position by skipping the start codon to produce peptides with the sequence XVDYKDDDDK (X = **7**, **13**-**15**). Following FLAG-tag enrichment, LC-HRMS confirmed initiation with monomers **7** and **13**-**15** (Fig. 6b). When the mass corresponding to the *m/z* = M+2H (**13-15**) or *m/z* = M+3H (**7**) charge state for each peptide was extracted from the total ion chromatogram, there was a clear peak for each peptide. No such peak was observed in reactions that lacked either the DNA template or acyl-tRNA, confirming templated ribosomal initiation with α-thio acid **13**, 2-benzylmalonic acid **14**, and *N*-formyl-L-α-amino acid **15**. Two peaks of identical mass are present in the EIC when translation was initiated with 2-benzylmalonic acid **14**, which we assign to diastereomeric peptides resulting from acylation at either the *pro*-S or *pro*-R carboxylate. Combined with the multiple peaks present in the RNAse A assay with 2-benzylmalonic acids **17, 18,** and **19** (Fig. 4a and Supplementary Fig. 7), as well as the two *meta*-CF_3_-2-BMA conformations observed in the structure of *Ma*FRSA (Fig. 5b), the two peptide products of identical mass generated in the IVT reactions imply that *Ma*FRSx enzymes can effectively acylate either of the two prochiral carboxylates of a malonic acid substrate.

**Figure 6.**
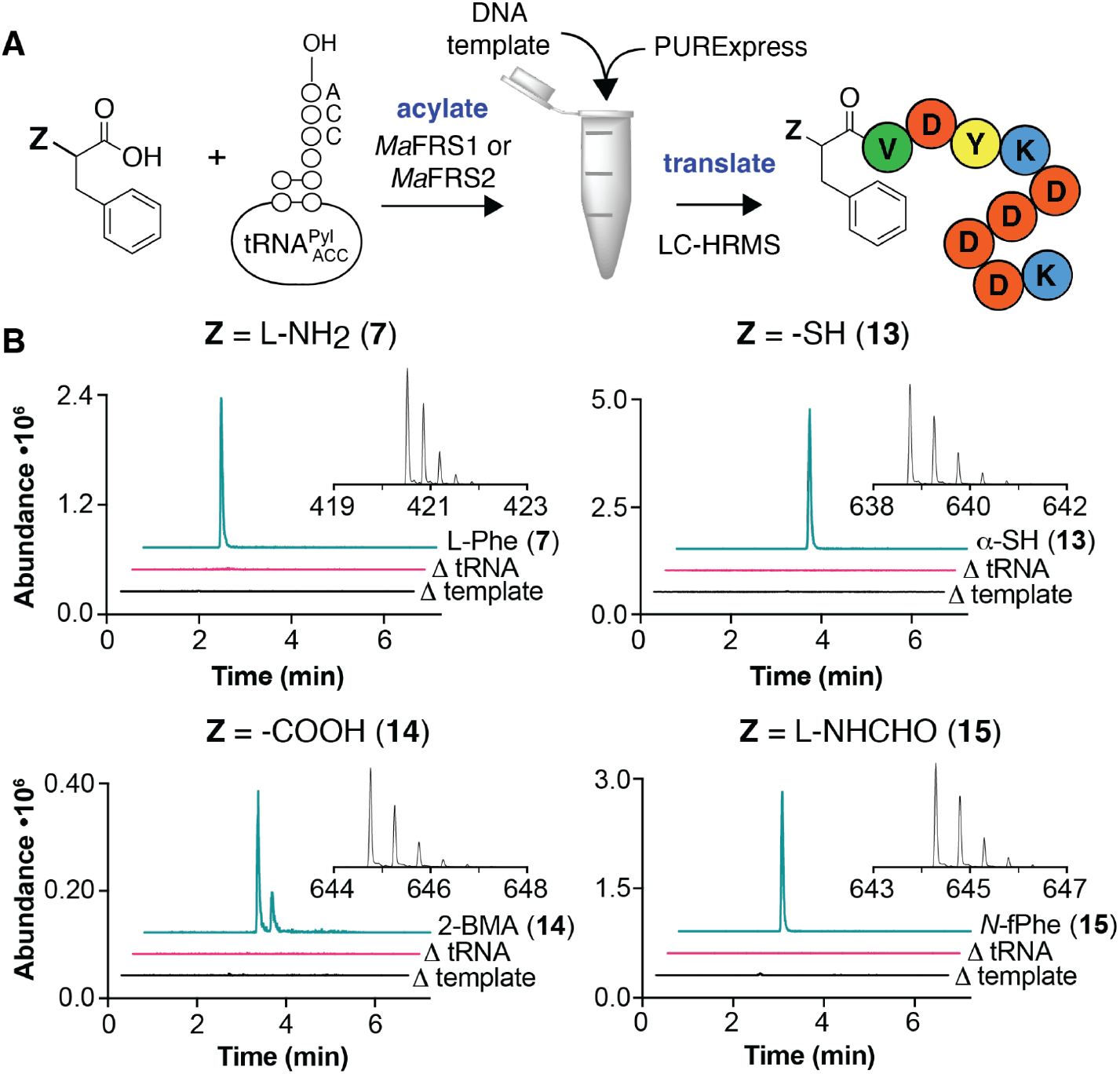
*In vitro* translation (IVT) of peptides initiated with novel monomers *via* codon skipping. **a**, Workflow for *in vitro* translation. **b**, Extracted ion chromatograms (EICs) and mass spectra of peptide products obtained using *Ma*-tRNA^Pyl^-ACC charged with monomers **7**, **13**-**15** by *Ma*FRS1 (**7** and **15**) or *Ma*FRS2 (**13** and **14**). Insets show mass spectra for major ions used to generate the EIC of the translated peptide initiated with the indicated monomer. Expected (exp) and observed (obs) *m/z* peaks in mass spectra are as follows: L-Phe **7** (M+3H), exp: 420.51906, obs: 420.52249; α-SH **13** (M+2H), exp: 638.75554, obs: 638.75614; *N*-fPhe **15** (M+2H), exp: 644.27242, obs: 644.27167; and 2-BMA **14** (M+2H), exp: 644.76442, obs: 644.76498.

## Discussion

Expanding and reprogramming the genetic code for the templated biosynthesis of sequence-defined hetero-polymers demands orthogonal aminoacyl-tRNA synthetase/tRNA pairs that accept non-L-α-amino acid substrates. Fahnestock & Rich reported over fifty years ago that the peptidyl transferase center (PTC) of the *E. coli* ribosome tolerates an α-hydroxyl substituent in place of the α-amine of phenylalanine and described the *in vitro* biosynthesis of a polyester^3^.

More recent studies have broadened the scope of wild-type PTC reactivity to include other nucleophilic heteroatoms in place of the α-amine^13–16^. However, with the exception of substrates carrying an α-hydroxyl substituent^19–21^, none of these non-L-α-amino acid monomers are substrates for any known orthogonal aminoacyl-tRNA synthetase. The challenge is that most aminoacyl-tRNA synthetases with known structures simultaneously engage both the substrate α-amine and α-carboxylate moieties via multiple main-chain and side-chain hydrogen bonds to position the latter for adenylation and acylation. These well-conserved hydrogen bond networks complicate the engineering of new enzymes that accept substrates with alternative amine/carboxylate orientations or conformations (such as β-amino acids or aramids), or those whose α-substituents are large and/or electrostatically distinctive (such as malonates, α-thio acids, and .*N*-acyl α-amino acids).

Yokoyama and coworkers demonstrated that the water-mediated hydrogen-bond network in the active site of *Methanosarcina mazei* pyrrolysyl-tRNA synthetase facilitated recognition of substrates with conservative substitutions (α-OH, α-H, α-NHCH_3_) of the α-amino group^20^. Here we expand the scope of monomers recognized and processed by PylRS variants to include α-thio acids and *N*-formyl-L-α-amino acids as well as those that carry an α-carboxyl functional group in place of the α-amine: prochiral malonic acids with protein-like side chains. Although thioesters and malonic acids are ubiquitous intermediates in polyketide and fatty acid biosynthesis^59,60^, as far as we know, tRNAs enzymatically acylated with a polyketide precursor or an α-thio acid represent novel chemical species. Such tRNAs could forge a new synthetic link between ribosomal translation and assembly-line polyketide synthases^61^, the two molecular machines responsible for protein and polyketide biosynthesis, respectively. The availability of multiple mutually orthogonal synthetases that accept α-hydroxy acids provides a route to the *in vivo* production of mixed sequence polyesters. Combined with synthetic genomes^62,63^ ribosomes capable of carbon-carbon bond formation would set the stage for the template-driven biosynthesis of unique hybrid biomaterials and sequence-defined polyketide-peptide oligomers, such as those produced by PKS-NRPS biosynthetic modules.

## Methods

### Expression and purification of *Ma*PylRS, *Ma*FRS1, *Ma*FRS2, and *Ma*FRSA for biochemistry

Plasmids used to express wild-type (WT) *Ma*PylRS (pET32a-*Ma*PylRS) and *Ma*FRS1 (pET32a-*Ma*FRS1) were constructed by inserting synthetic dsDNA fragments (Supplementary Table 1) into the NdeI-NdeI cut sites of a pET32a vector using the Gibson method^64^. pET32a-*Ma*FRS2 and pET32a-*Ma*FRSA were constructed from pET32a-*Ma*FRS1 using a Q5® Site-Directed Mutagenesis Kit (NEB). Primers pRF31 & pRF32, and pRF32 & pRF33 (Supplementary Table 1) were used to construct pET32a-*Ma*FRS2 and pET32a-*Ma*FRSA, respectively. The sequences of the plasmids spanning the inserted regions were confirmed via Sanger sequencing at the UC Berkeley DNA Sequencing Facility using primers T7 F and T7 R (Supplementary Table 1) and the complete sequence of each plasmid was confirmed by the Massachusetts General Hospital CCIB DNA Core.

Chemically competent cells were prepared by following a modified published protocol^65^. Briefly, 5 mL of LB was inoculated using a freezer stock of BL21-Gold (DE3)pLysS cells. The following day, 50 mL of LB was inoculated with 0.5 mL of the culture from the previous day and incubated at 37 °C with shaking at 200 rpm until the culture reached an OD_600_ between 0.3-0.4. The cells were collected by centrifugation at 4303 × g for 20 min at 4 °C. The cell pellet was resuspended in 5 mL of sterile filtered TSS solution (10% w/v polyethylene glycol 8000, 30 mM MgCl_2_, 5% v/v DMSO in 25 g/L LB). The chemically competent cells were portioned into 100 μL aliquots in 1.5 mL microcentrifuge tubes, flash frozen in liquid N_2_, and stored at −80 °C until use. The following protocol was used to transform plasmids into chemically competent cells: 20 μL of KCM solution (500 mM KCl, 150 mM CaCl_2_, 250 mM MgCl_2_) was added to a 100 μL aliquot of cells on ice along with approximately 200 ng of the requisite plasmid and water to a final volume of 200 μL. The cells were incubated on ice for 30 min and then heat-shocked by placing them for 90 s in a water-bath heated to 42 °C. Immediately after heat shock the cells were placed on ice for 2 min, after which 800 μL of LB was added. The cells then incubated at 37 °C with shaking at 200 rpm for 60 min. The cells were plated onto LB-agar plates with the appropriate antibiotic and incubated overnight at 37 °C.

Plasmids used to express wild type (WT) *Ma*PylRS, *Ma*FRS1, *Ma*FRS2 and *Ma*FRSA were transformed into BL21-Gold (DE3)pLysS chemically competent cells and plated onto LB agar plates supplemented with 100 μg/mL carbenicillin. Colonies were picked the following day and used to inoculate 10 mL of LB supplemented with 100 μg/mL carbenicillin. The cultures were incubated overnight at 37 °C with shaking at 200 rpm. The following day the 10 mL cultures were used to inoculate 1 L of LB supplemented with 100 μg/mL carbenicillin in 4 L baffled Erlenmeyer flasks. Cultures were incubated at 37 °C with shaking at 200 rpm for 3 h until they reached an OD600 of 0.6-0.8. At this point, isopropyl β-D-1-thiogalactopyranoside (IPTG) was added to a final concentration of 1 mM and incubation was continued for 6 h at 30 °C with shaking at 200 rpm. Cells were harvested by centrifugation at 4303 × g for 20 min at 4 °C and the cell pellets were stored at −80 °C until the expressed protein was purified as described below.

The following buffers were used for protein purification: Wash buffer: 50 mM sodium phosphate (pH 7.4), 500 mM NaCl, 20 mM β-mercaptoethanol (BME), 25 mM imidazole; Elution buffer: 50 mM sodium phosphate (pH 7.4), 500 mM NaCl, 20 mM β-mercaptoethanol (BME), 100 mM imidazole; Storage buffer: 100 mM HEPES-K, pH 7.2, 100 mM NaCl, 10 mM MgCl_2_, 4 mM dithiothreitol (DTT), 20% v/v glycerol. 1 cOmplete Mini EDTA-free protease inhibitor tablet was added to Wash and Elution buffers immediately before use. To isolate protein, cell pellets were resuspended in Wash buffer (5 mL/g cells). The resultant cell paste was lysed at 4 °C by sonication (Branson Sonifier 250) over 10 cycles consisting of 30 sec sonication followed by 30 sec manual swirling. The lysate was centrifuged at 4303 × g for 10 min at 4 °C to separate the soluble and insoluble fractions. The soluble lysate was incubated at 4 °C with 1 mL of Ni-NTA agarose resin (washed with water and equilibrated with Wash buffer) for 2 h. The lysate-resin mixture was added to a 65 g RediSep® Disposable Sample Load Cartridge (Teledyne ISCO) and allowed to drain at RT. The protein-bound Ni-NTA agarose resin was then washed with three 10 mL aliquots of Wash buffer. The protein was eluted from Ni-NTA agarose resin by rinsing the resin three times with 10 mL Elution buffer. The elution fractions were pooled and concentrated using a 10 kDa MWCO Amicon® Ultra-15 Centrifugal Filter Unit (4303 × g, 4 °C). The protein was then buffer-exchanged into Storage buffer until the [imidazole] was < 5 μM using the same centrifugal filter unit. The protein was dispensed into 20 μL single-use aliquots and stored at −80 °C for up to 8 months. Protein concentration was measured using the Bradford assay ^66^. Yields were between 8 and 12 mg/L. Proteins were analyzed by SDS-PAGE (Supplementary Fig. 2a) using Any kD™ Mini-PROTEAN® TGX™ Precast Protein Gels (BioRad). The gels were run at 200 V for 30 min.

Proteins were analyzed by LC-MS to confirm their identities (Supplementary Fig. 2b). Samples analyzed by mass spectrometry were resolved using a Poroshell StableBond 300 C8 (2.1 x 75 mm, 5 μm, Agilent Technologies part #660750-906) using a 1290 Infinity II UHPLC (G7120AR, Agilent). The mobile phases used for separation were (A) 0.1% formic acid in water and (B) 100% acetonitrile, and the flow rate was 0.4 mL/min. After an initial hold at 5% (B) for 0.5 min, proteins were eluted using a linear gradient from 5 to 75% (B) for 9.5 min, a linear gradient from 75 to 100% (B) for 1 min, a hold at 100% (B) for 1 min, a linear gradient 100 to 5% (B) for 3.5 min, and finally a hold at 5% (B) for 4.5 min. Protein masses were analyzed using LC-HRMS with an Agilent 6530 Q-TOF AJS-ESI (G6530BAR). The following parameters were used: gas temperature 300 °C, drying gas flow 12 L/min, nebulizer pressure 35 psi, sheath gas temperature 350 °C, sheath gas flow 11 L/min, fragmentor voltage 175 V, skimmer voltage 65 V, Oct 1 RF Vpp 750 V, Vcap 3500 V, nozzle voltage 1000 V, 3 spectra/s.

Analytical size exclusion chromatography (Supplementary Fig. 2C) was performed on an ÄKTA Pure 25. A flow rate of 0.5 mL/min was used for all steps. A Superdex® 75 Increase 10/300 GL column (stored and operated at 4 °C) was washed with 1.5 column volumes (CV) of degassed and sterile filtered MilliQ water. The column equilibrated in 1.5 column volumes of SEC Buffer: 100 mM HEPES (pH 7.2), 100 mM NaCl, 10 mM MgCl_2_, 4 mM DTT. Approximately 800 μg of protein in 250 μL SEC Buffer was loaded into a 500 μL capillary loop. The sample loop was washed with 2.0 mL of SEC Buffer as the sample was injected onto the column. The sample was eluted in 1.5 column volumes of SEC Buffer and analyzed by UV absorbance at 280 nm.

### Transcription and purification of tRNAs

The DNA template used for transcribing *M. αlvus* tRNA^Pyl^ (*Ma*-tRNA^Pyl^)^35^ was prepared by annealing and extending the ssDNA oligonucleotides *Ma*-PylT-F and *Ma*-PylT-R (2 mM, Supplementary Table 1) using OneTaq 2x Master Mix (NEB). The annealing and extension used the following protocol on a thermocycler (BioRad C1000 Touch™): 94 °C for 30 s, 30 cycles of [94 °C for 20 s, 53 °C for 30 s, 68 °C for 60 s], 68 ° C for 300 s. Following extension, the reaction mixture was supplemented with sodium acetate (pH 5.2) to a final concentration of 300 mM, washed once with 1:1 (*v*/*v*) acid phenol:chloroform, twice with chloroform, and the dsDNA product precipitated upon addition of ethanol to a final concentration of 71%. The pellet was resuspended in water and the concentration of dsDNA determined using a NanoDrop ND-1000 (Thermo Scientific). The template begins with a single C preceding the T7 promoter, which increases yields of T7 transcripts^67^. The penultimate residue of *Ma*-PylT-R carries a 2’-methoxy modification, which reduces non-templated nucleotide addition by T7 RNA polymerase during *in vitro* transcription^68^.

*Ma*-tRNA^Pyl^ was transcribed *in vitro* using a modified version of a published procedure^69^. Transcription reactions (25 μL) contained the following components: 40 mM Tris-HCl (pH 8.0), 100 mM NaCl, 20 mM DTT, 2 mM spermidine, 5 mM adenosine triphosphate (ATP), 5 mM cytidine triphosphate (CTP), 5 mM guanosine triphosphate (GTP), 5 mM uridine triphosphate (UTP), 20 mM guanosine monophosphate (GMP), 0.2 mg/mL bovine serum albumin, 20 mM MgCl_2_, 12.5 ng/μL DNA template, 0.025 mg/mL T7 RNA polymerase. These reactions were incubated at 37 °C in a thermocycler for 3 h. Four 25 μL reactions were pooled, and sodium acetate (pH 5.2) was added to a final concentration of 300 mM in a volume of 200 μL. The transcription reactions were extracted once with 1:1 (*v*/*v*) acid phenol:chloroform, washed twice with chloroform, and the tRNA product precipitated by adding ethanol to a final concentration of 71%. After precipitation, the tRNA pellet was resuspended in water and incubated with 8 U of RQ1 RNAse-free DNAse (Promega) at 37 °C for 30 min according to the manufacturer’s protocol. The tRNA was then washed with phenol:chloroform and chloroform as described above, precipitated, and resuspended in water. To remove small molecules, the tRNA was further purified using a Micro Bio-Spin™ P-30 Gel Column, Tris Buffer RNase-free (Bio-Rad) after first exchanging the column buffer to water according to the manufacturer’s protocol. The tRNA was precipitated once more, resuspended in water, quantified using a NanoDrop ND-1000, aliquoted, and stored at −20 °C.

tRNA was analyzed by Urea-PAGE (Supplementary Fig. 2d) using a 10% Mini-PROTEAN® TBE-Urea Gel (BioRad). The gels were run at 120 V for 30 min then stained with SYBR-Safe gel stain (Thermo-Fisher) for 5 minutes before imaging. *Ma*-tRNA^Pyl^ was analyzed by LC-MS to confirm its identity. Samples were resolved on a ACQUITY UPLC BEH C18 Column (130 Å, 1.7 μm, 2.1 mm X 50 mm, Waters part # 186002350, 60 °C) using an ACQUITY UPLC I-Class PLUS (Waters part # 186015082). The mobile phases used were (A) 8 mM triethylamine (TEA), 80 mM hexafluoroisopropanol (HFIP), 5 μM ethylenediaminetetraacetic acid (EDTA, free acid) in 100% MilliQ water; and (B) 4 mM TEA, 40 mM HFIP, 5 μM EDTA (free acid) in 50% MilliQ water/50% methanol. The method used a flow rate of 0.3 mL/min and began with Mobile Phase B at 22% that increased linearly to 40 % B over 10 min, followed by a linear gradient from 40 to 60% B for 1 min, a hold at 60% B for 1 min, a linear gradient from 60 to 22% B over 0.1 min, then a hold at 22% B for 2.9 min. The mass of the RNA was analyzed using LC-HRMS with a Waters Xevo G2-XS Tof (Waters part #186010532) in negative ion mode with the following parameters: capillary voltage: 2000 V, sampling cone: 40, source offset: 40, source temperature: 140 °C, desolvation temperature: 20 °C, cone gas flow: 10 L/h, desolvation gas flow: 800 L/h, 1 spectrum/s. Expected masses of oligonucleotide products were calculated using the AAT Bioquest RNA Molecular Weight Calculator ^70^. Deconvoluted mass spectra were obtained using the MaxEnt software (Waters Corporation).

### Procedure for RNAse A assays

Reaction mixtures (25 μL) used to acylate tRNA contained the following components: 100 mM Hepes-K (pH 7.5), 4 mM DTT, 10 mM MgCl_2_, 10 mM ATP, 0 - 10 mM substrate, 0.1 U *E. coli* inorganic pyrophosphatase (NEB), 25 μM *Ma*-tRNA^Pyl^, and 2.5 μM enzyme (*Ma*PylRS, *Ma*FRS1, *Ma*FRS2, or *Ma*FRSA). Reaction mixtures were incubated at 37 °C in a dry-air incubator for 2 h. tRNA samples from enzymatic acylation reactions were quenched with 27.5 μL of RNAse A solution (1.5 U/μL RNAse A (Millipore-Sigma), 200 mM sodium acetate, pH 5.2) and incubated for 5 min at room temperature. Proteins were then precipitated upon addition of 50% trichloroacetic acid (TCA, Sigma-Aldrich) to a final concentration of 5%. After precipitating protein at −80 °C for 30 min, insoluble material was removed by centrifugation at 21,300 × g for 10 min at 4 °C. The soluble fraction was then transferred to autosampler vials, kept on ice until immediately before LC-MS analysis, and returned to ice immediately afterwards.

Samples analyzed by mass spectrometry were resolved using a Zorbax Eclipse XDB-C18 RRHD column (2.1 × 50 mm, 1.8 μm, room temperature, Agilent Technologies part # 981757-902) fitted with a guard column (Zorbax Eclipse XDB-C18, 2.1 x 5 mm 1.8 μm, Agilent part # 821725-903) using a 1290 Infinity II UHPLC (G7120AR, Agilent). The mobile phases used were (A) 0.1% formic acid in water; and (B) 100% acetonitrile. The method used a flow rate of 0.7 mL/min and began with Mobile Phase B held at 4% for 1.35 min, followed by a linear gradient from 4 to 40% B over 1.25 min, a linear gradient from 40 to 100% B over 0.4 min, a linear gradient from 100 to 4% B over 0.7 min, then finally B held at 4% for 0.8 min. Acylation was confirmed by correctly identifying the exact mass of the 2’ and 3’ acyl-adenosine product corresponding to the substrate tested in the extracted ion chromatogram by LC-HRMS with an Agilent 6530 Q-TOF AJS-ESI (G6530BAR). The following parameters were used: fragmentor voltage of 175 V, gas temperature of 300°C, gas flow of 12 L/min, sheath gas temperature of 350°C, sheath gas flow of 12 L/min, nebulizer pressure of 35 psi, skimmer voltage of 65 V, Vcap of 3500 V, and collection rate of 3 spectra/s. Expected exact masses of acyl-adenosine nucleosides (Supplementary Table 2) were calculated using ChemDraw 19.0 and extracted from the total ion chromatograms ± 100 ppm.

### Procedure for determining aminoacylation yields using intact tRNA mass spectrometry

Enzymatic tRNA acylation reactions (25 μL) were performed as described in **III**. Sodium acetate (pH 5.2) was added to the acylation reactions to a final concentration of 300 mM in a volume of 200 μL. The reactions were then extracted once with a 1:1 (v/v) mixture of acidic phenol (pH 4.5):chloroform and washed twice with chloroform. After extraction, the acylated tRNA was precipitated by adding ethanol to a final concentration of 71% and incubation at −80 °C for 30 min, followed by centrifugation at 21,300 × g for 30 min at 4 °C. After the supernatant was removed, acylated tRNA was resuspended in water and kept on ice for analysis.

tRNA samples from enzymatic acylation reactions were analyzed by LC-MS as described in **II**. Because the unacylated tRNA peak in each total ion chromatogram (TIC) contains tRNA species that cannot be enzymatically acylated (primarily tRNAs that lack the 3’ terminal adenosine^71^), simple integration of the acylated and non-acylated peaks in the A_260_ chromatogram does not accurately quantify the acylation yield. To accurately quantify acylation yield, we used the following procedure. For each sample, the mass data was collected between 500 and 2000 *m/z*. A subset of the mass data collected defined as the raw MS deconvolution range (panel C in Supplementary Fig. 3 - 6) was used to produce the deconvoluted mass spectra (panel D in Supplementary Fig. 3 - 6). The raw MS deconvolution range of each macromolecule species contains multiple peaks that correspond to different charge states of that macromolecule. Within the raw mass spectrum deconvolution range we identified the most abundant charge state peak in the raw mass spectrum of each tRNA species (unacylated tRNA, monoacylated tRNA, and diacylated tRNA), which is identified as the major ion by an asterisk in panel C in figs. S3 to S6. To quantify the relative abundance of each species, the exact mass of the major ions ± 0.3000 Da was extracted from the TIC to produce extracted ion chromatograms (EICs, panel E in Supplementary Fig. 3 - 6). The EICs were integrated and the areas of the peaks that aligned with the correct peaks in the TIC (as determined from the deconvoluted mass spectrum) were used for quantification of yields (Supplementary Table 3). For malonic acid substrates, the integrated peak areas for the EICs from both the malonic acid product and the decarboxylation product are added together to determine the overall acylation yield. Each sample was injected 3 times; chromatograms and spectra in Supplementary Fig. 3 - 6 are representative, yields shown in Supplementary Table 3 are an average of the 3 injections. Expected masses of oligonucleotide products were calculated using the AAT Bioquest RNA Molecular Weight Calculator^70^ and the molecular weights of the small molecules added to them were calculated using ChemDraw 19.0. All masses identified in the mass spectra are summarized in Data S1.

### Malachite green assay to monitor adenylation

Enzymatic adenylation reactions were monitored using malachite green using a previous protocol with modifications^49^. Each adenylation reaction (60 μL) contained the following components: 200 mM HEPES-K (pH 7.5), 4 mM DTT, 10 mM MgCl_2_, 0.2 mM ATP, 0 - 10 mM substrate, 4 U/mL *E. coli* inorganic pyrophosphatase (NEB), and 2.5 μM enzyme (*Ma*FRS1 or *Ma*FRSA). Adenylation reactions were incubated at 37 °C in a dry-air incubator. Aliquots (10 μL) were withdrawn after 0, 5, 10, 20, and 30 min and quenched upon addition to an equal volume of 20 mM EDTA (pH 8.0) on ice. Once all aliquots were withdrawn, 80 μL of Malachite Green Solution (Echelon Biosciences) was added to each aliquot and the mixture incubated at RT for 30 min. After shaking for 30 sec to remove bubbles, the absorbance at 620 nm was measured on a Synergy HTX plate reader (BioTek). The absorbance was then converted to phosphate concentration using a phosphate standard curve (0 - 100 μM) and plotted over time to determine turnover numbers.

### Structure determination

The following synthetic dsDNA sequence was cloned upstream of *Ma*FRSA (*Ma*PylRS N166A:V168A) into pET32a-*Ma*FRSA by Gibson assembly ^64^ and used for subsequent crystallographic studies: GSS linker - 6xHis - SSG linker - thrombin site - *Ma*FRSA (Supplementary Table 1). The sequence of the pET32a-6xHis-thrombin-MaFRSA plasmid was confirmed with Sanger sequencing from Genewiz using primers T7 F and T7 R (Supplementary Table 1). The procedure used to express and purify *Ma*FRSA for crystallography using pET32a-6xHis-thrombin-*Ma*FRSA was adapted from a reported protocol used to express and purify wildtype *M. alvus* PylRS by Seki *et al*. ^54^. BL21(DE3) Gold competent cells (Agilent Technologies) were transformed with pET32a-6xHis-thrombin-*Ma*FRSA and grown in TB media at 37 °C. Protein expression was induced at an OD_600_ reading of 1.2 with 1 mM isopropyl β-D-1-thiogalactopyranoside (IPTG). The temperature was lowered to 20 °C and growth continued overnight. Cells were pelleted for 1 h at 4,300 × g and resuspended in Lysis buffer (50 mM potassium phosphate (pH 7.4), 25 mM imidazole, 500 mM sodium chloride, 5 mM β-mercaptoethanol, 1 cOmplete Mini EDTA-free protease inhibitor tablet). Cells were lysed by homogenization (Avestin Emulsiflex C3). After centrifugation for 1 h at 10,000 × g, the clarified lysate was bound to TALON® Metal Affinity Resin (Takara Bio) for 1 h at 4 °C, washed with additional lysis buffer, and eluted with Elution Buffer (50 mM potassium phosphate (pH 7.4), 500 mM imidazole, 500 mM sodium chloride, 5 mM β-mercaptoethanol). The eluate was dialyzed overnight at 4 °C into Cleavage buffer (40 mM potassium phosphate (pH 7.4), 100 mM NaCl, 1 mM dithiothreitol (DTT)) then incubated overnight at room temperature with thrombin protease on a solid agarose support (MilliporeSigma). Following cleavage, the protein was passed over additional TALON® resin to remove the 6xHis tag and dialyzed overnight at 4 °C into Sizing buffer (30 mM potassium phosphate (pH 7.4), 200 mM NaCl, 1 mM DTT). The protein was concentrated and loaded onto a HiLoad® 16/600 Superdex® 200 pg column (Cytiva Life Sciences) equilibrated with Sizing buffer on an ÄKTA Pure 25 fast-liquid chromatography machine. Purified *Ma*FRSA was dialyzed into Storage buffer (10 mM Tris-HCl (pH 8.0), 150 mM NaCl, 10 mM MgCl2, 10 mM β-mercaptoethanol), concentrated to 20 mg/mL, aliquoted, and flash-frozen for crystallography (Supplementary Fig. 2, F to G).

Initial crystallization screening conditions were adapted from Seki *et al*. ^54^. Crystals were grown by hanging drop vapor-diffusion in 24-well plates. 25 μL of 100 mM *meta*-trifluoromethyl-2-benzylmalonate (*meta*-CF_3_-2-BMA) pH ~7 was added to 1.5 mL micrcentrifuge tubes and the water was removed by evaporation. The dried aliquots were then resuspended at a concentration of 100 mM with *Ma*FRSA in Storage buffer at three concentrations (6.9, 12.3, and 19.2 mg/mL) and 10 mM adenosine 5’-(β,γ-imido)triphosphate lithium salt hydrate (AMP-PNP). The protein/substrate solution (1 μL) was mixed in a 1:1 ratio with the reservoir solution (1 μL) containing 10 mM Tris-HCl pH 7.4 and 26% polyethylene glycol 3350 and incubated over 1 mL of reservoir solution at 18 °C. Crystals with an octahedral shape appeared within one week. Crystals were plunged into liquid nitrogen to freeze with no cryoprotectant used.

Data were collected at the Advanced Light Source beamline 8.3.1 at 100 K with a wavelength of 1.11583 Å. Data collection and refinement statistics are presented in Supplementary Table 4. Diffraction data were indexed and integrated with XDS ^72^, then merged and scaled with Pointless ^73^ and Aimless ^74^. The crystals were in the space group I4, and the unit cell dimensions were 108.958, 108.958, and 112.26 Å. The structure was solved by molecular replacement with Phaser ^75^ using a single chain of the wild-type apo structure of *M. alvus* PylRS (PDB code: 6JP2) ^54^ as the search model. There were two copies of *Ma*FRSA in the asymmetric unit. The model was improved with iterative cycles of manual model building in COOT ^76^ alternating with refinement in Phenix ^77,78^ using data up to 1.8 Å resolution. Structural analysis and figures were generated using Pymol version 2.4.2 ^79^.

### *In vitro* translation initiation

The *Ma*-tRNA^Pyl^-ACC dsDNA template was prepared as described in **II** using the primers *Ma*-PylT-ACC F and *Ma*-PylT-ACC R (Supplementary Table 1). *Ma*-tRNA^Pyl^-ACC was also transcribed, purified, and analyzed as described previously. Enzymatic tRNA acylation reactions (150 μL) were performed as described in **III** with slight modifications. The enzyme concentration was increased to 12.5 μM (monomers **7**, **14**, and **15**) or 25 μM (monomer **13**) and the incubation time was increased to 3 hours at 37 °C. Sodium acetate (pH 5.2) was added to the acylation reactions to a final concentration of 300 mM in a volume of 200 μL. The reactions were then extracted once with a 1:1 (v/v) mixture of acidic phenol (pH 4.5):chloroform and washed twice with chloroform. After extraction, the acylated tRNA was precipitated by adding ethanol to a final concentration of 71% and incubation at −80 °C for 30 min followed by centrifugation at 21,300 × g for 30 min at 4 °C. Acylated tRNAs were resuspended in water to a concentration of 307 μM immediately before *in vitro* translation.

Templates for expression of MGVDYKDDDDK were prepared by annealing and extending the oligonucleotides MGVflag-1 and MGVflag-2 using Q5® High-Fidelity 2X Master Mix (NEB) (Supplementary Table 1). The annealing and extension used the following protocol on a thermocycler (BioRad C1000 Touch™): 98 °C for 30 s, 10 cycles of [98 °C for 10 s, 55 °C for 30 s, 72 °C for 45 s], 10 cycles of [98 °C for 10 s, 67 °C for 30 s, 72 °C for 45 s], and 72 ° C for 300 s. Following extension, the reaction mixture was supplemented with sodium acetate (pH 5.2) to a final concentration of 300 mM, extracted once with a 1:1 (*v*/*v*) mixture of basic phenol (pH 8.0):chloroform, and washed twice with chloroform. The dsDNA product was precipitated upon addition of ethanol to a final concentration of 71% and incubation at −80 °C for 30 min followed by centrifugation at 21,300 × g for 30 min at 4 °C. The dsDNA pellets were washed once with 70% (*v/v*) ethanol and resuspended in 10 mM Tris-HCl pH 8.0 to a concentration of 500 ng/μL and stored at −20 °C until use in translation.

*In vitro* transcription/translation by codon skipping of the short FLAG tag-containing peptides X-Val-Asp-Tyr-Lys-Asp-Asp-Asp-Asp-Lys (XV-Flag) where X = **7**, **13**, **14**, or **15** was carried out using the PURExpress® Δ (aa, tRNA) Kit (New England Biolabs, E6840S) based on a previous protocol with slight modifications ^16^. The XV-Flag peptides were produced with the following reactions (12.5 μL): Solution A (ΔtRNA, Δaa; 2.5 μL), amino acid stock mix (1.25 μL; 33 mM L-valine, 33 mM L-aspartic acid, 33 mM L-tyrosine, 33 mM L-lysine), tRNA solution (1.25 μL), Solution B (3.75 μL), 250 ng dsDNA MGVDYKDDDDK template (0.5 μL), and *Ma*-tRNA^Pyl^-ACC acylated with **7**, **13**, **14**, or **15** (3.25 μL). The reactions were incubated in a thermocycler (BioRad C1000 Touch™) at 37 °C for 2 hours and quenched by placement on ice.

Translated peptides were purified from *in vitro* translation reactions by enrichment using Anti-FLAG^®^ M2 Magnetic Beads (Millipore Sigma) according to the manufacturer’s protocol with slight modifications. For each peptide, 10 μL of a 50% (*v/v*) suspension of magnetic beads was used. The supernatant was pipetted from the beads on a magnetic manifold. The beads were then washed twice by incubating with 100 μL of TBS (150 mM NaCl, 50 mM Tris-HCl, pH 7.6) for 10 min at room temperature then removing the supernatant each time with a magnetic manifold. The *in vitro* translation reactions were added to the beads and incubated at RT for 30 min with periodic agitation. The beads were washed again three times with 100 μL of TBS as described above. Peptides were eluted by incubation with 12.5 μL of 0.1 M glycine-HCl pH 2.8 for 10 minutes. The supernatant was transferred to vials and kept on ice for analysis.

The purified peptides were analyzed based on a previous protocol ^16^. The supernatant was analyzed on an ZORBAX Eclipse XDB-C18 column (1.8 μm, 2.1 × 50 mm, room temperature, Agilent) using an 1290 Infinity II UHPLC (G7120AR, Agilent). The following method was used for separation: an initial hold at 95% Solvent A (0.1% formic acid in water) and 5% Solvent B (acetonitrile) for 0.5 min followed by a linear gradient from 5 to 50% Solvent B over 4.5 min at flow rate of 0.7 mL/min. Peptides were identified using LC-HRMS with an Agilent 6530 Q-TOF AJS-ESI (G6230BAR). The following parameters were used: a fragmentor voltage of 175 V, gas temperature of 300 °C, gas flow rate of 12 L/min, sheath gas temperature of 350 °C, sheath gas flow rate of 11 L/min, nebulizer pressure of 35 psi, skimmer voltage of 75 V, Vcap of 3500 V, and collection rate of 3 spectra/s. Expected exact masses of the major charge state for each peptide were calculated using ChemDraw 19.0 and extracted from the total ion chromatograms ± 100 ppm.

## Acknowledgements

We thank Drs. Hasan Celik and Alicia Lund, and UC Berkeley’s NMR facility in the College of Chemistry (CoC-NMR) for spectroscopic assistance, Prof. John Kuriyan for generously sharing crystallography equipment, Drs. James Holton and George Meigs for X-ray diffraction assistance, Dr. Shuai Zheng for advice on synthesis, and Isaac Knudson for advice on *in vitro* translation. The authors are grateful to Drs. Scott Miller, Jamie Cate, and Abhishek Chatterjee for meaningful discussion and feedback. This work was supported by the NSF Center for Genetically Encoded Materials (C-GEM), CHE 2002182. R.F. was supported by Agilent Technologies. C.S. was supported by a Berkeley Fellowship for Graduate Study. L.T.R. was supported by the NSF Graduate Research Fellowship Program under Grant No. 1752814. Beamline 8.3.1 at the Advanced Light Source is operated by the University of California at San Francisco with generous grants from the National Institutes of Health (R01 GM124149 and P30 GM124169), Plexxikon Inc., and the Integrated Diffraction Analysis Technologies program of the US Department of Energy Office of Biological and Environmental Research. The Advanced Light Source (Berkeley, CA) is a national user facility operated by Lawrence Berkeley National Laboratory on behalf of the US Department of Energy under contract number DE-AC02-05CH11231, Office of Basic Energy Sciences.

## Author contributions

R.F., C.S., O.A., and A.S. designed the project. R.F. led acylation experiments and assisted with crystallography and *in vitro* translation. C.S. led crystallography and *in vitro* translation and assisted with acylation experiments. L.T.R. synthesized materials and assisted with crystallography. N.H. assisted with acylation experiments. S.S. and C.G. assisted with crystallography. R.F., C.S., and A.S. prepared the manuscript.

## Additional Information

**Supplementary Information**

**Supplementary Figs. 1-10, materials, synthesis notes, references.**

**Supplementary Data.** Structural model of *Ma*FRSA is available in the Protein Data Bank with code 7U0R.

## References

1. Lutz, J., Ouchi, M., Liu, D. & Sawamoto, M. Sequence-Controlled Polymers. Science 341, 628–+ (2013).

2. Barnes, J. et al. Iterative exponential growth of stereo- and sequence-controlled polymers. Nat. Chem. 7, 810–815 (2015).

3. Fahnestock, S. & Rich, A. Ribosome-catalyzed polyester formation. Science 173, 340–343 (1971).

4. Ohta, A., Murakami, H., Higashimura, E. & Suga, H. Synthesis of polyester by means of genetic code reprogramming. Chem. Biol. 14, 1315–1322 (2007).

5. Katoh, T., Iwane, Y. & Suga, H. Logical engineering of D-arm and T-stem of tRNA that enhances d-amino acid incorporation. Nucleic Acids Res. 45, 12601–12610 (2017).

6. Fujino, T., Goto, Y., Suga, H. & Murakami, H. Ribosomal Synthesis of Peptides with Multiple β-Amino Acids. J. Am. Chem. Soc. 138, 1962–1969 (2016).

7. Katoh, T. & Suga, H. Ribosomal Incorporation of Consecutive β-Amino Acids. J. Am. Chem. Soc. 140, 12159–12167 (2018).

8. Adaligil, E., Song, A., Hallenbeck, K. K., Cunningham, C. N. & Fairbrother, W. J. Ribosomal Synthesis of Macrocyclic Peptides with β ^2^ - and β ^2,3^-Homo-Amino Acids for the Development of Natural Product-Like Combinatorial Libraries. ACS Chem. Biol. 16, 1011–1018 (2021).

9. Katoh, T., Sengoku, T., Hirata, K., Ogata, K. & Suga, H. Ribosomal synthesis and de novo discovery of bioactive foldamer peptides containing cyclic β-amino acids. Nat. Chem. 12, 1081–1088 (2020).

10. Katoh, T. & Suga, H. Ribosomal Elongation of Cyclic γ-Amino Acids using a Reprogrammed Genetic Code. J. Am. Chem. Soc. 142, 4965–4969 (2020).

11. Adaligil, E., Song, A., Cunningham, C. N. & Fairbrother, W. J. Ribosomal Synthesis of Macrocyclic Peptides with Linear γ ^4^ - and β-Hydroxy-γ ^4^-amino Acids. ACS Chem. Biol. 16, 1325–1331 (2021).

12. Lee, J., Schwarz, K. J., Kim, D. S., Moore, J. S. & Jewett, M. C. Ribosome-mediated polymerization of long chain carbon and cyclic amino acids into peptides in vitro. Nat. Commun. 11, 4304 (2020).

13. Katoh, T. & Suga, H. Consecutive Ribosomal Incorporation of α-Aminoxy/α-Hydrazino Acids with l / d -Configurations into Nascent Peptide Chains. J. Am. Chem. Soc. 143, 18844–18848 (2021).

14. Takatsuji, R. et al. Ribosomal Synthesis of Backbone-Cyclic Peptides Compatible with In Vitro Display. J. Am. Chem. Soc. 141, 2279–2287 (2019).

15. Katoh, T. & Suga, H. Ribosomal Elongation of Aminobenzoic Acid Derivatives. J. Am. Chem. Soc. 142, 16518–16522 (2020).

16. Ad, O. et al. Translation of Diverse Aramid- and 1,3-Dicarbonyl-peptides by Wild Type Ribosomes *in Vitro*. ACS Cent. Sci. 5, 1289–1294 (2019).

17. Sievers, A., Beringer, M., Rodnina, M. V. & Wolfenden, R. The ribosome as an entropy trap. Proc. Natl. Acad. Sci. U. S. A. 101, 7897 (2004).

18. England, P. M., Zhang, Y., Dougherty, D. A. & Lester, H. A. Backbone Mutations in Transmembrane Domains of a Ligand-Gated Ion Channel: Implications for the Mechanism of Gating. Cell 96, 89–98 (1999).

19. Guo, J., Wang, J., Anderson, J. C. & Schultz, P. G. Addition of an α-Hydroxy Acid to the Genetic Code of Bacteria. Angew. Chem. 120, 734–737 (2008).

20. Kobayashi, T., Yanagisawa, T., Sakamoto, K. & Yokoyama, S. Recognition of Non-α-amino Substrates by Pyrrolysyl-tRNA Synthetase. J. Mol. Biol. 385, 1352–1360 (2009).

21. Li, Y.-M. et al. Ligation of Expressed Protein α-Hydrazides *via* Genetic Incorporation of an α-Hydroxy Acid. ACS Chem. Biol. 7, 1015–1022 (2012).

22. Melo Czekster, C., Robertson, W. E., Walker, A. S., Söll, D. & Schepartz, A. In Vivo Biosynthesis of a β-Amino Acid-Containing Protein. J. Am. Chem. Soc. 138, 5194–5197 (2016).

23. Chen, S., Ji, X., Gao, M., Dedkova, L. M. & Hecht, S. M. In Cellulo Synthesis of Proteins Containing a Fluorescent Oxazole Amino Acid. J. Am. Chem. Soc. 141, 5597–5601 (2019).

24. Liu, C. C. & Schultz, P. G. Adding New Chemistries to the Genetic Code. Annu. Rev. Biochem. 79, 413–444 (2010).

25. Wan, W., Tharp, J. M. & Liu, W. R. Pyrrolysyl-tRNA synthetase: An ordinary enzyme but an outstanding genetic code expansion tool. Biochim. Biophys. Acta BBA - Proteins Proteomics 1844, 1059–1070 (2014).

26. Vargas-Rodriguez, O., Sevostyanova, A., Söll, D. & Crnković, A. Upgrading aminoacyl-tRNA synthetases for genetic code expansion. Curr. Opin. Chem. Biol. 46, 115–122 (2018).

27. Italia, J. S. et al. Mutually Orthogonal Nonsense-Suppression Systems and Conjugation Chemistries for Precise Protein Labeling at up to Three Distinct Sites. J. Am. Chem. Soc. 141, 6204–6212 (2019).

28. Chin, J. W. Expanding and reprogramming the genetic code. Nature 550, 53–60 (2017).

29. Srinivasan Gayathri, James Carey M., & Krzycki Joseph A. Pyrrolysine Encoded by UAG in Archaea: Charging of a UAG-Decoding Specialized tRNA. Science 296, 1459–1462 (2002).

30. Ambrogelly, A., Palioura, S. & Soll, D. Natural expansion of the genetic code. Nat. Chem. Biol. 3, 29–35 (2007).

31. Wang, L., Brock, A., Herberich, B. & Schultz, P. G. Expanding the Genetic Code of *Escherichia coli*. Science 292, 498–500 (2001).

32. Kobayashi, T. et al. Structural basis for orthogonal tRNA specificities of tyrosyl-tRNA synthetases for genetic code expansion. Nat. Struct. Biol. 10, 8 (2003).

33. Yanagisawa, T. et al. Crystallographic Studies on Multiple Conformational States of Activesite Loops in Pyrrolysyl-tRNA Synthetase. J. Mol. Biol. 378, 634–652 (2008).

34. Borrel, G. et al. Genome Sequence of ‘Candidatus Methanomethylophilus alvus’ Mx1201, a Methanogenic Archaeon from the Human Gut Belonging to a Seventh Order of Methanogens. J. Bacteriol. 194, 6944–6945 (2012).

35. Willis, J. C. W. & Chin, J. W. Mutually orthogonal pyrrolysyl-tRNA synthetase/tRNA pairs. Nat. Chem. 10, 831–837 (2018).

36. Wang, Y.-S. et al. The de novo engineering of pyrrolysyl-tRNA synthetase for genetic incorporation of l-phenylalanine and its derivatives. Mol. Biosyst. 7, 714 (2011).

37. Wang, Y.-S., Fang, X., Wallace, A. L., Wu, B. & Liu, W. R. A Rationally Designed Pyrrolysyl-tRNA Synthetase Mutant with a Broad Substrate Spectrum. J. Am. Chem. Soc. 134, 2950–2953 (2012).

38. Herring, S. et al. The amino-terminal domain of pyrrolysyl-tRNA synthetase is dispensable in vitro but required for in vivo activity. FEBS Lett. 581, 3197–3203 (2007).

39. McMurry, J. L. & Chang, M. C. Y. Fluorothreonyl-tRNA deacylase prevents mistranslation in the organofluorine producer *Streptomyces cattleya*. Proc. Natl. Acad. Sci. 114, 11920–11925 (2017).

40. Findly, D., Herries, D. G., Mathias, A. P., Rabin, B. R. & Ross, C. A. The Active Site and Mechanism of Action of Bovine Pancreatic Ribonuclease. Nature 190, 781–784 (1961).

41. Griffin, B. E., Jarman, M., Reese, C. B., Sulston, J. E. & Trentham, D. R. Some Observations Relating to Acyl Mobility in Aminoacyl Soluble Ribonucleic Acids*. Biochemistry 5, 3638–3649 (1966).

42. Stepanov, V. G., Moor, N. A., Ankilova, V. N. & Lavrik, O. I. Phenylalanyl-tRNA synthetase from *Thermus thermophilus* can attach two molecules of phenylalanine to tRNA ^Phe^. FEBS Lett. 311, 192–194 (1992).

43. Wang, B., Zhou, J., Lodder, M., Anderson, R. D. & Hecht, S. M. Tandemly Activated tRNAs as Participants in Protein Synthesis. J. Biol. Chem. 281, 13865–13868 (2006).

44. Ko, J. et al. Pyrrolysyl-tRNA synthetase variants reveal ancestral aminoacylation function. FEBS Lett. 587, 3243–3248 (2013).

45. Xuan, W. et al. Site-Specific Incorporation of a Thioester Containing Amino Acid into Proteins. ACS Chem. Biol. 13, 578–581 (2018).

46. Hwang, S., Lee, N., Cho, S., Palsson, B. & Cho, B.-K. Repurposing Modular Polyketide Synthases and Non-ribosomal Peptide Synthetases for Novel Chemical Biosynthesis. Front. Mol. Biosci. 7, (2020).

47. Varshney, U., Lee, C. P., Seong, B. L. & RajBhandary, U. L. Mutants of initiator tRNA that function both as initiators and elongators. J. Biol. Chem. 266, 18018–18024 (1991).

48. Tharp, J. M. et al. Initiation of Protein Synthesis with Non-Canonical Amino Acids In Vivo. Angew. Chem. Int. Ed. 59, 3122–3126 (2020).

49. Cestari, I. & Stuart, K. A Spectrophotometric Assay for Quantitative Measurement of Aminoacyl-tRNA Synthetase Activity. J. Biomol. Screen. 18, 490–497 (2013).

50. Wang, Y.-S. et al. Genetic Incorporation of Twelve *meta*-Substituted Phenylalanine Derivatives Using a Single Pyrrolysyl-tRNA Synthetase Mutant. ACS Chem. Biol. 8, 405–415 (2013).

51. Lee, Y.-J. et al. Genetically encoded fluorophenylalanines enable insights into the recognition of lysine trimethylation by an epigenetic reader. Chem. Commun. 52, 12606–12609 (2016).

52. Guo, L.-T. et al. Polyspecific pyrrolysyl-tRNA synthetases from directed evolution. Proc. Natl. Acad. Sci. 111, 16724–16729 (2014).

53. Tharp, J. M., Wang, Y.-S., Lee, Y.-J., Yang, Y. & Liu, W. R. Genetic Incorporation of Seven *ortho* -Substituted Phenylalanine Derivatives. ACS Chem. Biol. 9, 884–890 (2014).

54. Seki, E., Yanagisawa, T., Kuratani, M., Sakamoto, K. & Yokoyama, S. Fully Productive Cell-Free Genetic Code Expansion by Structure-Based Engineering of *Methanomethylophilus alvus* Pyrrolysyl-tRNA Synthetase. ACS Synth. Biol. 9, 718–732 (2020).

55. Kavran, J. M. et al. Structure of pyrrolysyl-tRNA synthetase, an archaeal enzyme for genetic code innovation. Proc. Natl. Acad. Sci. 104, 11268–11273 (2007).

56. Englert, M. et al. Probing the active site tryptophan of *Staphylococcus aureus* thioredoxin with an analog. Nucleic Acids Res. 43, 11061–11067 (2015).

57. Laursen, B. S., Sørensen, H. P., Mortensen, K. K. & Sperling-Petersen, H. U. Initiation of Protein Synthesis in Bacteria. Microbiol. Mol. Biol. Rev. 69, 101–123 (2005).

58. Lee, J. et al. Expanding the limits of the second genetic code with ribozymes. Nat. Commun. 10, 5097 (2019).

59. Nivina, A., Yuet, K., Hsu, J. & Khosla, C. Evolution and Diversity of Assembly-Line Polyketide Synthases. Chem. Rev. 119, 12524–12547 (2019).

60. Tsai, S. The Structural Enzymology of Iterative Aromatic Polyketide Synthases: A Critical Comparison with Fatty Acid Synthases. in Annual Review of Biochemistry (ed. Kornberg, R.) vol. 87 503–531 (2018).

61. Walsh, C. T., O’Brien, R. V. & Khosla, C. Nonproteinogenic amino acid building blocks for nonribosomal peptide and hybrid polyketide scaffolds. Angew. Chem. Int. Ed. 52, 7098–7124 (2013).

62. Ostrov, N. et al. Design, synthesis, and testing toward a 57-codon genome. Science 353, 819–822 (2016).

63. Fredens, J. et al. Total synthesis of Escherichia coli with a recoded genome. Nature 569, 514–518 (2019).

64. Gibson, D. G. et al. Enzymatic assembly of DNA molecules up to several hundred kilobases. Nat. Methods 6, 343–345 (2009).

65. Chung, C. T., Niemela, S. L. & Miller, R. H. One-step preparation of competent Escherichia coli: transformation and storage of bacterial cells in the same solution. Proc. Natl. Acad. Sci. 86, 2172–2175 (1989).

66. Bradford, M. M. A Rapid and Sensitive Method for the Quantitation of Microgram Quantities of Protein Utilizing the Principle of Protein-Dye Binding. 7.

67. Baklanov, M. Effect on DNA transcription of nucleotide sequences upstream to T7 promoter. Nucleic Acids Res. 24, 3659–3660 (1996).

68. Kao, C., Zheng, M. & Rüdisser, S. A simple and efficient method to reduce nontemplated nucleotide addition at the 3’ terminus of RNAs transcribed by T7 RNA polymerase. RNA 5, 1268–1272 (1999).

69. Noncanonical Amino Acids: Methods and Protocols. vol. 1728 (Springer New York, 2018).

70. RNA Molecular Weight Calculator | AAT Bioquest. https://www.aatbio.com/tools/calculate-RNA-molecular-weight-mw/.

71. Sprinzl, M. & Cramer, F. The -C-C-A End of tRNA and Its Role in Protein Biosynthesis. in Progress in Nucleic Acid Research and Molecular Biology vol. 22 1–69 (Elsevier, 1979).

72. Kabsch, W. XDS. Acta Crystallogr. D Biol. Crystallogr. 66, 125–132 (2010).

73. Evans, P. R. An introduction to data reduction: space-group determination, scaling and intensity statistics. Acta Crystallogr. D Biol. Crystallogr. 67, 282–292 (2011).

74. Evans, P. R. & Murshudov, G. N. How good are my data and what is the resolution? Acta Crystallogr. D Biol. Crystallogr. 69, 1204–1214 (2013).

75. Bunkóczi, G. et al. Phaser.MRage: automated molecular replacement. Acta Crystallogr. D Biol. Crystallogr. 69, 2276–2286 (2013).

76. Emsley, P., Lohkamp, B., Scott, W. G. & Cowtan, K. Features and development of Coot. Acta Crystallogr. D Biol. Crystallogr. 66, 486–501 (2010).

77. Adams, P. D. et al. PHENIX: a comprehensive Python-based system for macromolecular structure solution. Acta Crystallogr. D Biol. Crystallogr. 66, 213–221 (2010).

78. Liebschner, D. et al. Macromolecular structure determination using X-rays, neutrons and electrons: recent developments in Phenix. Acta Crystallogr. Sect. Struct. Biol. 75, 861–877 (2019).

79. Schrödinger, LLC. The PyMOL Molecular Graphics System, Version 1.8. (2015).

80. Herold, S., Bafaluy, D. & Muñiz, K. Anodic benzylic C(sp3)–H amination: unified access to pyrrolidines and piperidines. Green Chem. 20, 3191–3196 (2018).

81. Matveeva, E. D. et al. Syntheses of Compounds Active toward Glutamate Receptors: II. Synthesis of Spiro Hydantoins of the Indan Series. Russ. J. Org. Chem. 38, 1769–1774 (2002).

82. Bindman, N. A., Bobeica, S. C., Liu, W. R. & van der Donk, W. A. Facile Removal of Leader Peptides from Lanthipeptides by Incorporation of a Hydroxy Acid. J. Am. Chem. Soc. 137, 6975–6978 (2015).

83. Madeira, F. et al. The EMBL-EBI search and sequence analysis tools APIs in 2019. Nucleic Acids Res. 47, W636–W641 (2019).

